# A live-attenuated SARS-CoV-2 vaccine candidate with accessory protein deletions

**DOI:** 10.1101/2022.02.14.480460

**Authors:** Yang Liu, Xianwen Zhang, Jianying Liu, Hongjie Xia, Jing Zou, Antonio E. Muruato, Sivakumar Periasamy, Jessica A. Plante, Nathen E. Bopp, Chaitanya Kurhade, Alexander Bukreyev, Ping Ren, Tian Wang, Menachery Vineet D., Kenneth S. Plante, Xuping Xie, Scott C. Weaver, Pei-Yong Shi

**Affiliations:** Department of Biochemistry and Molecular Biology, University of Texas Medical Branch, Galveston TX, USA; Department of Microbiology and Immunology, University of Texas Medical Branch, Galveston TX, USA; Institute for Human Infections and Immunity, University of Texas Medical Branch, Galveston, TX, USA; Department of Pathology, University of Texas Medical Branch, Galveston, TX, USA; Galveston National Laboratory, Galveston, Texas, USA; World Reference Center for Emerging Viruses and Arboviruses, University of Texas Medical Branch, Galveston, TX, USA; Sealy Institute for Vaccine Sciences, University of Texas Medical Branch, Galveston, TX, USA; Sealy Institute for Drug Discovery, University of Texas Medical Branch, Galveston, TX, USA; Sealy Center for Structural Biology & Molecular Biophysics, University of Texas Medical Branch, Galveston, TX, USA

**Author notes:** Correspondence: K.S.P., X.X., S.C.W., or P.-Y.S. These authors contributed equally (Yang Liu, Xianwen Zhang, Jianying Liu, Hongjie Xia). Lead contact: P.-Y.S.

**Keywords:** SARS-CoV-2, COVID-19, coronavirus, diagnosis, vaccine, antiviral, variant

## Abstract

We report a live-attenuated SARS-CoV-2 vaccine candidate with (i) re-engineered viral transcriptional regulator sequences and (ii) deleted open-reading-frames (ORF) 3, 6, 7, and 8 (Δ3678). The Δ3678 virus replicates about 7,500-fold lower than wild-type SARS-CoV-2 on primary human airway cultures, but restores its replication on interferon-deficient Vero-E6 cells that are approved for vaccine production. The Δ3678 virus is highly attenuated in both hamster and K18-hACE2 mouse models. A single-dose immunization of the Δ3678 virus protects hamsters from wild-type virus challenge and transmission. Among the deleted ORFs in the Δ3678 virus, ORF3a accounts for the most attenuation through antagonizing STAT1 phosphorylation during type-I interferon signaling. We also developed an mNeonGreen reporter Δ3678 virus for high-throughput neutralization and antiviral testing. Altogether, the results suggest that Δ3678 SARS-CoV-2 may serve as a live-attenuated vaccine candidate and a research tool for potential biosafety level-2 use.

## Introduction

The pandemic of COVID-19, caused by SARS-CoV-2, has led to over 395 million confirmed infections and 5.7 million deaths (as of February 6, 2022; https://coronavirus.jhu.edu/). Different vaccine platforms have been successfully developed for COVID-19 at an unprecedented pace, including mRNA, viral vector, subunit protein, and inactivated virus. Live-attenuated vaccines of SARS-CoV-2 have not been actively explored, even though they may have advantages of low cost, strong immunogenicity, and long immune durability. The SARS-CoV-2 virion consists of an internal nucleocapsid, formed by the genomic RNA coated with nucleocapsid (N) proteins, and an external envelope, formed by a cell-derived bilipid membrane embedded with spike (S), membrane (M), and envelope (E) proteins^1^. The plus-sense, single-stranded viral RNA genome encodes open-reading-frames (ORFs) for replicase (ORF1a/ORF1b), S, E, M, and N structural proteins, as well as seven additional ORFs for accessory proteins^2^. Although the exact functions of SARS-CoV-2 accessory proteins remain to be determined, previous studies of other coronaviruses suggest that these proteins are not essential for viral replication but can modulate replication and pathogenesis through interacting with host pathways^3–7^. Thus, deletion of the accessory proteins could be used to attenuate SARS-CoV-2.

Reverse genetic systems are important tools to engineer and study viruses. In response to the COVID-19 pandemic, three types of reverse genetic systems have been developed for SARS-CoV-2: (i) an infectious cDNA clone^8–12^, (ii) a transient replicon (a self-replicating viral RNA with one or more genes deleted)^13, 14^, and (iii) a *trans*-complementation system (replicon RNAs in cells that express the missing genes in the replicon, allowing for single cycle replication without spread)^15–17^. The three systems have their own strengths and weaknesses and are complementary to each other when applied to address different research questions. The infectious cDNA clone requires biosafety level-3 (BSL-3) containment to recover and handle infectious SARS-CoV-2. The transient replicon system requires RNA preparation and transfection for each experiment; cell lines harboring replicons that can be continuously cultured, like those developed for hepatitis C virus and other plus-sense RNA viruses^18–20^, have yet to be established for SARS-CoV-2. The *trans*-complementation system produces virions that can infect naïve cells for only a single round. Compared with the infectious cDNA clone, both the replicon and *trans*-complementation system have the advantage of allowing experiments to be performed at biosafety level-2 (BSL-2). A new system that combines the strengths of the current three systems (*e.g.*, multiple rounds of viral infection of naïve cells that can be performed at BSL2) would be very useful for COVID-19 research and countermeasure development. Here we report a highly attenuated SARS-CoV-2 (with deleted accessory proteins and rewired transcriptional regulator sequences) that can potentially serve as a live-attenuated vaccine platform and a BSL-2 experimental system.

## Results

### Attenuation of SARS-CoV-2 by deletion of accessory genes

Using an infectious clone of USA-WA1/2020 SARS-CoV-2, we constructed two mutant viruses containing accessory ORF deletions (**Fig. 1a**), one with ORF 6, 7, and 8 deletions (Δ678) and another with ORF 3, 6, 7, and 8 deletions (Δ3678). Besides the ORF deletions, the viral transcription regulatory sequences (TRS) of both Δ678 and Δ3678 viruses were mutated from the wild-type (WT) ACGAAC to <u>C</u>CG<u>G</u>A<u>T</u> (mutant nucleotides underlined; **Fig. 1a**). The mutated TRS virtually eliminates the possibility to produce WT SARS-CoV-2 through recombination between the Δ678 or Δ3678 RNA and inadvertently contaminating viral RNA^21, 22^. On Vero-E6 cells, the Δ678 virus developed plaques similar to the WT virus, whereas the Δ3678 virus produced smaller plaques (**Fig. 1b**). Both Δ678 and Δ3678 viruses were visible under the negative staining electron microscope (**Extended Data Fig. 1**). Replication kinetics analysis showed that WT and Δ678 replicated to comparable viral titers on Vero-E6 (**Fig. 1c**), Calu-3 (**Fig. 1d**), and primary human airway epithelial (HAE) cultures (**Fig. 1e**). In contrast, the replication of Δ3678 was slightly attenuated on Vero-E6 cells (**Fig. 1c**), but became significantly more attenuated on Calu-3 (360-fold lower peak viral titer than WT virus at 72 h; **Fig. 1d**) and HAE cultures (7,500-fold lower peak viral titer than WT virus on day 6; **Fig. 1e**). Consistently, the intracellular level of Δ3678 RNA was about 100-fold lower than that of Δ678 RNA and WT in HAE cells (**Fig. 1f**). Corroboratively, the Δ3678 virus caused much fewer cytopathic effects (CPE) than the Δ678 and WT viruses on both Vero-E6 and Calu-3 cells (**Extended Data Fig. 2**). To further confirm the attenuation of Δ3678 virus, we engineered the mNeonGreen (mNG) gene into the Δ3678 and WT viruses^23^. When infecting HAE cultures, the mNG Δ3678 virus developed significantly fewer mNG-positive cells than the mNG WT virus (**Fig. 1g**). Taken together, the results indicate that (i) deletions of ORFs 6, 7, and 8 slightly attenuate SARS-CoV-2 in cell culture; (ii) an additional deletion of ORF3 to the Δ678 virus significantly increases the attenuation of Δ3678; and (iii) the Δ3678 virus is strikingly more attenuated when infecting immune-competent cells than when infecting interferon-deficient cells.

**Figure 1.**
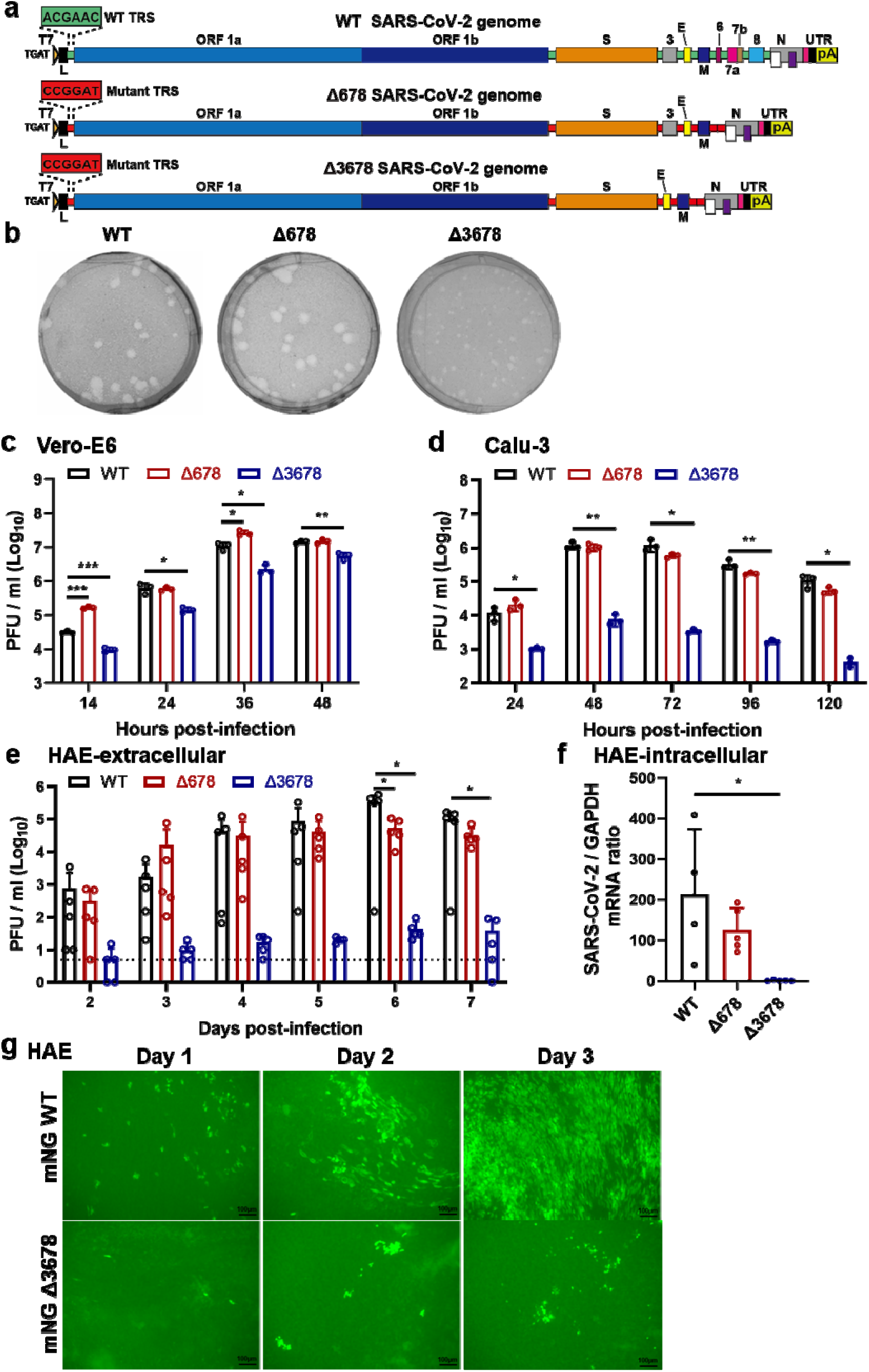
Attenuation of Δ3678 SARS-CoV-2 in cell culture. **a,** Scheme diagram for the construction of Δ678 and Δ3678 SARS-CoV-2. The deletions were introduced to the backbone of USA-WA1/2020 strain. T7, T7 promoter; L, leader sequence; TRS, transcription regulatory sequences; ORF, open reading frame; E, envelope glycoprotein gene; M, membrane glycoprotein gene; N, nucleocapsid gene; UTR, untranslated region; pA, poly A tails. **b,** Plaque morphologies of recombinant WT, Δ678, and Δ3678 viruses. Plaque assays were performed on Vero-E6 cells and stained on day 2.5 post-infection. **c-e,** Replication kinetics of WT, Δ678, and Δ3678 SARS-CoV-2s on Vero-E6 (**c)**, Calu-3 (**d)**, and HAE (**e)** cells. WT, Δ678, and Δ3678 viruses were inoculated onto Vero-E6, Calu-3, and HAE cells at MOIs of 0.01, 0.1, and 2, respectively. After a 2-h incubation, the cells were washed three times with DPBS and continuously cultured under fresh 2% FBS DMEM. Culture supernatants were measured for infectious virus titers using plaque assays on Vero-E6 cells. **f,** Intracellular levels of WT, Δ678, and Δ3678 RNA in HAE cells on day 7 post-infection. The HAE cells were washed with DPBS for three times, lysed by Trizol for RNA isolation, quantified for viral RNAs using RT-qPCR. Dots represent individual biological replicates (n=3 for Vero-E6 and Calu-3; n=5 for HAE). The values in the graph represent the mean ± standard deviation. An unpaired two-tailed *t* test was used to determine significant differences between WT and Δ678/Δ3678 groups. *P* values were adjusted using the Bonferroni correction to account for multiple comparisons. Differences were considered significant if p<0.025; P<0.025, *; P<0.005, **; and P<0.0005, ***. **g,** mNG-positive HAE cells after infection with mNG WT or mNG Δ3678 virus at an MOI of 0.5. Scale bar, 100 µm.

### Characterization of Δ3678 SARS-CoV-2 as a potential live-attenuated vaccine in a hamster model

We characterized the attenuation of Δ3678 virus in a hamster model (**Fig. 2a**). After intranasal infection with 10^6^ plaque-forming units (PFU) of Δ3678, the hamsters did not lose weight (**Fig. 2b**) or develop disease (**Fig. 2c**). In contrast, the WT virus-infected animals lost weight (**Fig. 2b**) and developed disease (**Fig. 2c**), as observed in our previous studies^24–26^. On day 2 post-infection, viral loads in the Δ3678-infected nasal washes (**Fig. 2d**), oral swabs (**Fig. 2e**), tracheae, and lungs (**Fig. 2f**) were 180-, 20-, 16-, and 100-fold lower than those in the WT-infected specimens. The Δ3678 infection elicited robust neutralization with a peak 50% neutralization titer (NT_50_) of 1,090 on day 21 post-infection, while the WT virus evoked 1.4-fold higher peak NT_50_ (**Fig. 2g**). The results demonstrate that the Δ3678 virus is attenuated and can elicit robust neutralization in hamsters.

**Figure 2.**
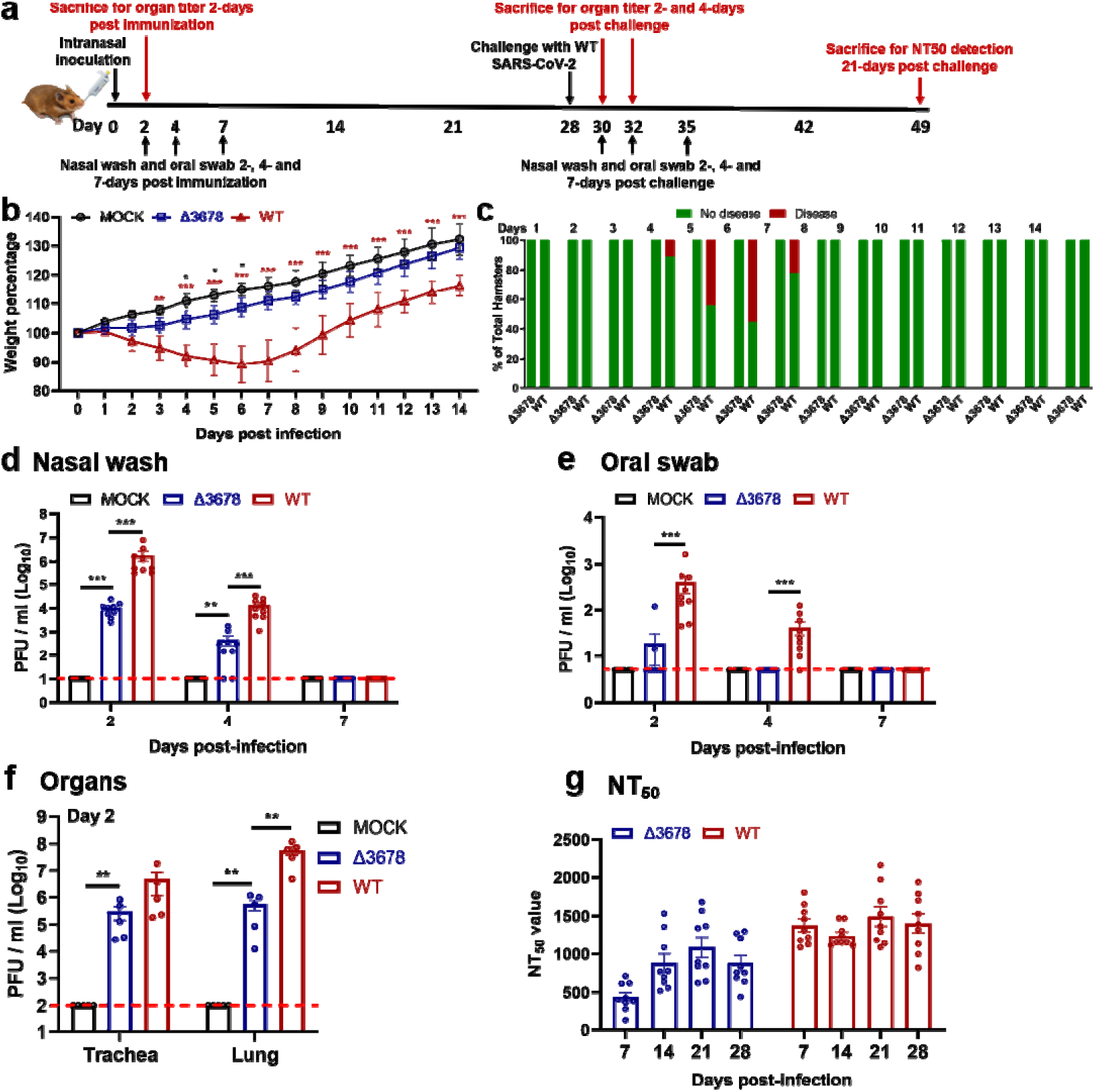
Attenuation of Δ3678 SARS-CoV-2 in hamsters. **a,** Experimental scheme of Δ3678 virus immunization and WT virus challenge. Hamsters were intranasally (I.N.) inoculated with 10^6^ PFU of WT or Δ3678 virus. On day 2 post-inoculation, organ viral loads (n=5) were measured by plaque assays on Vero-E6 cells. Nasal washes and oral swabs (n=10) were collected on days 2, 4, and 7 post-inoculation. On day 28 post-immunization, the hamsters were challenged by 10^5^ PFU of WT SARS-CoV-2. On days 2 and 4 post-challenge, plaque assays were performed to measure organ viral loads (n=5). On day 21 post-challenge, the animals were terminated to measure neutralization titer (NT_50_). **b,** Weight changes of hamsters after intranasal infection with WT (n=9) or Δ3678 (n=9) SARS-CoV-2. Uninfected mock group (n=9) was included as a negative control. Body weights were measured daily for 14 days. The data are shown as mean ± standard deviation. The weight changes between Δ3678 and mock or WT groups were analyzed using two-factor analysis of variance (ANOVA) with Tukey’s post hoc test. The black and red asterisks stand for the statistic difference between Δ3678 and mock or WT, respectively. *, P<0.05; **, P<0.01; ***, P<0.001. **c,** Disease of Δ678 and Δ3678 virus-infected animals. The diseases include ruffled fur, lethargic, hunched posture, and orbital tightening. The percentages of animals with or without diseases are presented. **d-f,** Viral loads in nasal wash (**d)**, oral swab (**e),** trachea, and lung (**f**) after infection with Δ3678 or WT virus. Dots represent individual animals (n=5). The mean ± standard error is presented. A non-parametric two-tailed Mann-Whitney test was used to determine the differences between mock, Δ3678, or WT groups. *P* values were adjusted using the Bonferroni correction to account for multiple comparisons. Differences were considered significant if p<0.025. *, P<0.025; **, P<0.005; ***, P<0.0005. **g,** Neutralization titers of sera from WT- and Δ3678 virus-inoculated hamsters on days 7, 14, 21, and 28 post-inoculation. The neutralization titers were measured against WT SARS-CoV-2.

We examined whether the above immunized hamsters could be protected from SARS-CoV-2 challenge. After intranasal challenge with 10^5^ PFU of WT SARS-CoV-2 on day 28 post-immunization (**Fig. 2a**), both the Δ3678- and WT virus-immunized animals were protected from weight loss (**Fig. 3a**) or disease (**Fig. 3b**). Compared with the mock-immunized group, the viral loads in the nasal washes (**Fig. 3c**) and oral swabs (**Fig. 3d**) from the Δ3678- and WT virus-immunized groups were decreased by >660 (day 2) and >80 folds (day 2), respectively; no infectious viruses were detected in trachea (**Fig. 3e**) and lungs (**Fig. 3f**) from the immunized groups. The challenge significantly increased the neutralization titers (on day 21 post-challenge) in both the Δ3678- and WT virus-immunized groups (**Fig. 3g**), suggesting that a single infection with the Δ3678 or WT virus did not elicit sterilizing immunity. Histopathology analysis showed that immunization with attenuated Δ3678 virus reduced lung pathology score, inflammation, alveolar septa change, and airway damage (**Extended Data Fig. 3**). In contrast, previous infection with WT virus did not exhibit improved lung histopathology after the challenge, possibly because the observed pathologic changes were caused by the initial WT viral infection (**Extended Data Fig. 3**). Collectively, the results demonstrate that immunization with attenuated Δ3678 virus can protect against WT SARS-CoV-2 challenge in hamsters.

**Figure 3.**
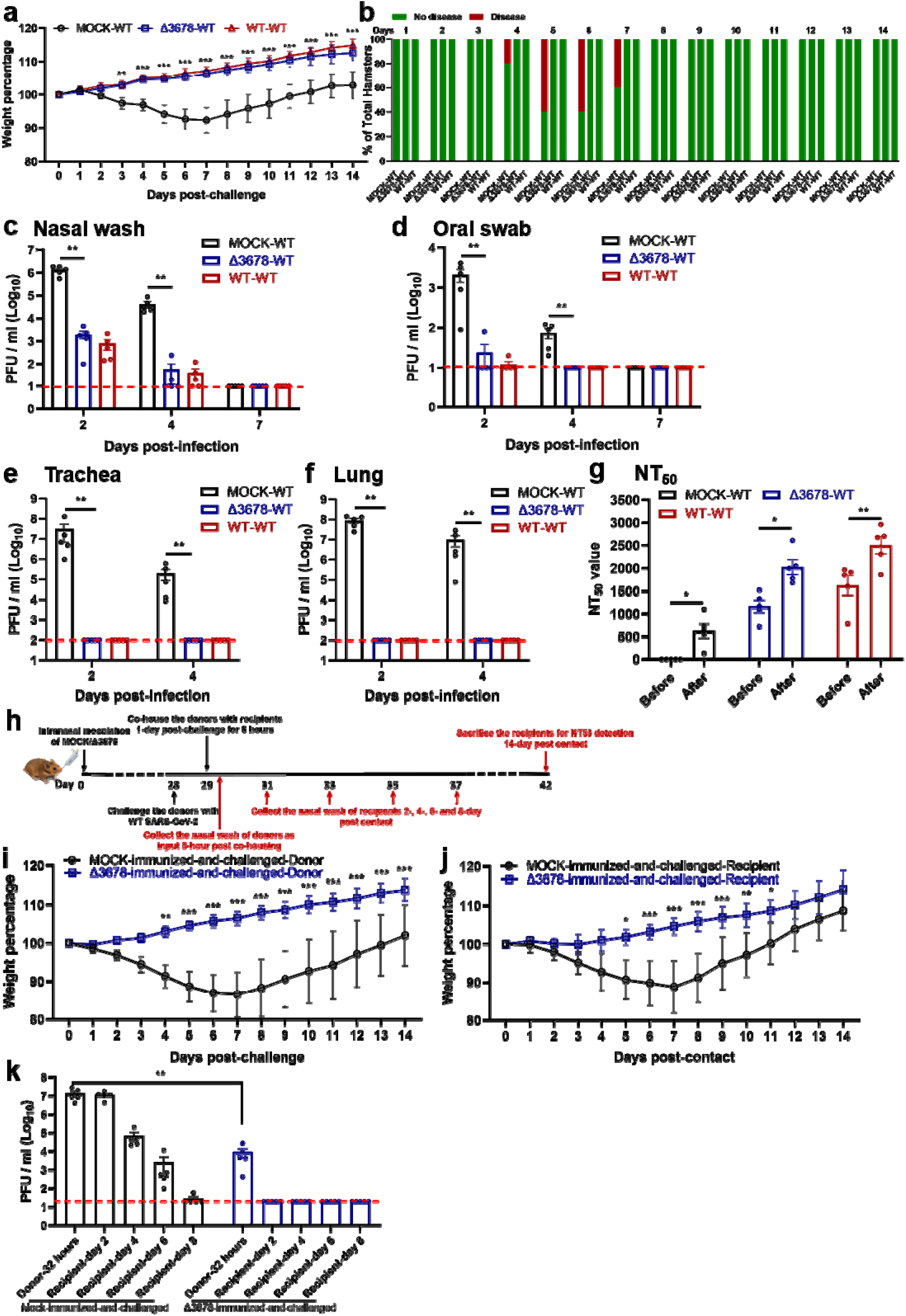
Protection of Δ3678 virus-immunized hamsters from WT SARS-CoV-2 challenge and transmission. **a, b,** Weight loss (**a**) and diseases (**b**) of immunized and challenged hamsters. **a,** Mock-immunized (n=5), Δ3678 virus-immunized (n=5), and WT virus-inoculated (n=5) hamsters were challenged with 10^5^PFU of WT SARS-CoV-2. The body weights were measured daily for 14 days post-challenge. The data are shown as mean ± standard deviation. The weight changes between Δ3678- and mock- or WT-inoculated groups were statistically analyzed using two-factor analysis of variance (ANOVA) with Tukey’s post hoc test. No statistical difference was observed between the Δ3678- and WT-inoculated groups. The statistic difference between the Δ3678- and mock-immunized groups are indicated. **, P<0.01; ***, P<0.001. **b,** After the challenge, animals developed diseases, including ruffled fur, lethargic, hunched posture, and orbital tightening. The percentages of animals with or without diseases were presented. **c-f,** Viral loads in the nasal wash (**c**), oral swab (**d**), trachea (**e**), and lung (**f**) after challenge. Dots represent individual animals (n=5). The values of mean ± standard error of the mean are presented. A non-parametric two-tailed Mann-Whitney test was used to determine the statistical differences between Δ3678-immunized and mock-immunized or WT-inoculated groups. *P* values were adjusted using the Bonferroni correction to account for multiple comparisons. Differences were considered significant if p<0.025. *, P<0.025; **, P<0.005. **g,** Neutralization titers of immunized hamsters before and after challenge. The “before challenge” sera were collected on day 28 post-immunization. The “after challenge” sera were collected on day 21 post-challenge. **h,** Experimental design of transmission blockage in hamsters. Hamsters were immunized with 10^5^PFU of Δ3678 virus (n=5) or medium mock (n=5). In day 28 post-immunization, the hamsters were challenged with 10^4^ PFU of WT SARS-CoV-2; these animals served as transmission donors. On day 1 post-challenge, the donor hamsters were co-housed with clean recipient hamsters for 8 h. The nasal washes of donor hamsters were collected immediately after contact (*i.e.*, 32 h post-challenge). The nasal washes of recipient hamsters were collected on days 2, 4, 6, and 8 post-contact. **i-j,** Weight loss of donors post-challenge (**i**) and recipients post-contact (**j**). The data are shown as mean ± standard deviation. The weight changes were statistically analyzed using two-factor analysis of variance (ANOVA) with Tukey’s post hoc test. *, P<0.05; **, P<0.01; ***, P<0.001. **k,** Viral loads in nasal wash of donors post-challenge and recipients post-contact. Dots represent individual animals (n=5). The values in the graph represent the mean ± standard error of mean. A non-parametric two-tailed Mann-Whitney test was used to analyze the difference between the mock-immunized-and-challenged and Δ3678-immunized-and-challenged hamsters. **, P<0.01.

Next, we tested whether lower dose immunization could also achieve protection. Hamsters were immunized with 10^2^, 10^3^, 10^4^, or 10^5^ PFU of Δ3678 virus. No weight loss was observed for all dose groups (**Extended Data Fig. 4a**). All dose groups developed equivalent, low lung viral loads (**Extended Data Fig. 4b**). After challenge with WT SARS-CoV-2, all dose groups exhibited protection similar to the 10^6^-PFU-dose group, including undectectable viral loads in the tracheae or lungs and significantly reduced viral loads in the nasal washes and oral swabs (**Extended Data Fig. 4c**). These results indicate a low dose of 10^2^ PFU of Δ3678 immunization is protective in hamsters.

Since infectious viruses were detected in the nasal and oral specimens from the Δ3678- immunized hamsters after the challenge, we examined whether such low levels of virus could be transmitted to naïve hamsters. On day 1 post-challenge, the Δ3678-immunized-and- challenged animals (donor) were co-housed with clean naive hamsters (recipient) for 8 h, after which the donor and recipient animals were separated (**Fig. 3h**). As expected, after WT virus challenge, no weight loss was observed in the Δ3678-immunized donor animals, but weight loss did occur in the mock-immunized donor animals (**Fig. 3i**). After co-housing with Δ3678- immunized-and-challenged donor animals, naïve recipient animals did not lose weight (**Fig. 3j**) and did not have infectious viruses in the nasal washes (**Fig. 3k**). In contrast, after co-housing with the mock-immunized-and-challenged donor animals, the recipient animals lost weight (**Fig. 3j**) and developed high viral loads in the nasal wash (**Fig. 3k**). Altogether, the results indicate that although the Δ3678-immunized donor animals developed low viral loads in their nasal and oral specimens after the WT virus challenge, they were unable to transmit the virus to clean naïve hamsters.

### Attenuation of Δ3678 SARS-CoV-2 in K18-hACE2 mice

To further characterize the attenuation of Δ3678, we intranasally inoculated K18-hACE2 mice with 4, 40, 400, 4,000, or 40,000 PFU of WT or Δ3678 virus (**Fig. 4a**). The infected groups were compared for their weight loss, survival rates, and disease signs. The WT viral infection caused weight loss at doses of ≥400 PFU (**Fig. 4b**), diseases at dose ≥400 PFU (**Fig. 4d**), and deaths at doses ≥4,000 PFU (**Fig. 4f**). In contrast, the Δ3678 virus caused slight (statistically insignificant) weight loss at 40,000 PFU (**Fig. 4c**), transient disease at ≥4,000 PFU (**Fig. 4e**), and no death at any dose (**Fig. 4g**). Consistently, the lung viral loads from the Δ3678-infected mice were significantly lower than those from the WT-infected animals (**Fig. 4h**). The results demonstrated that Δ3678 virus was highly attenuated in the K18-hACE2 mice.

**Figure 4.**
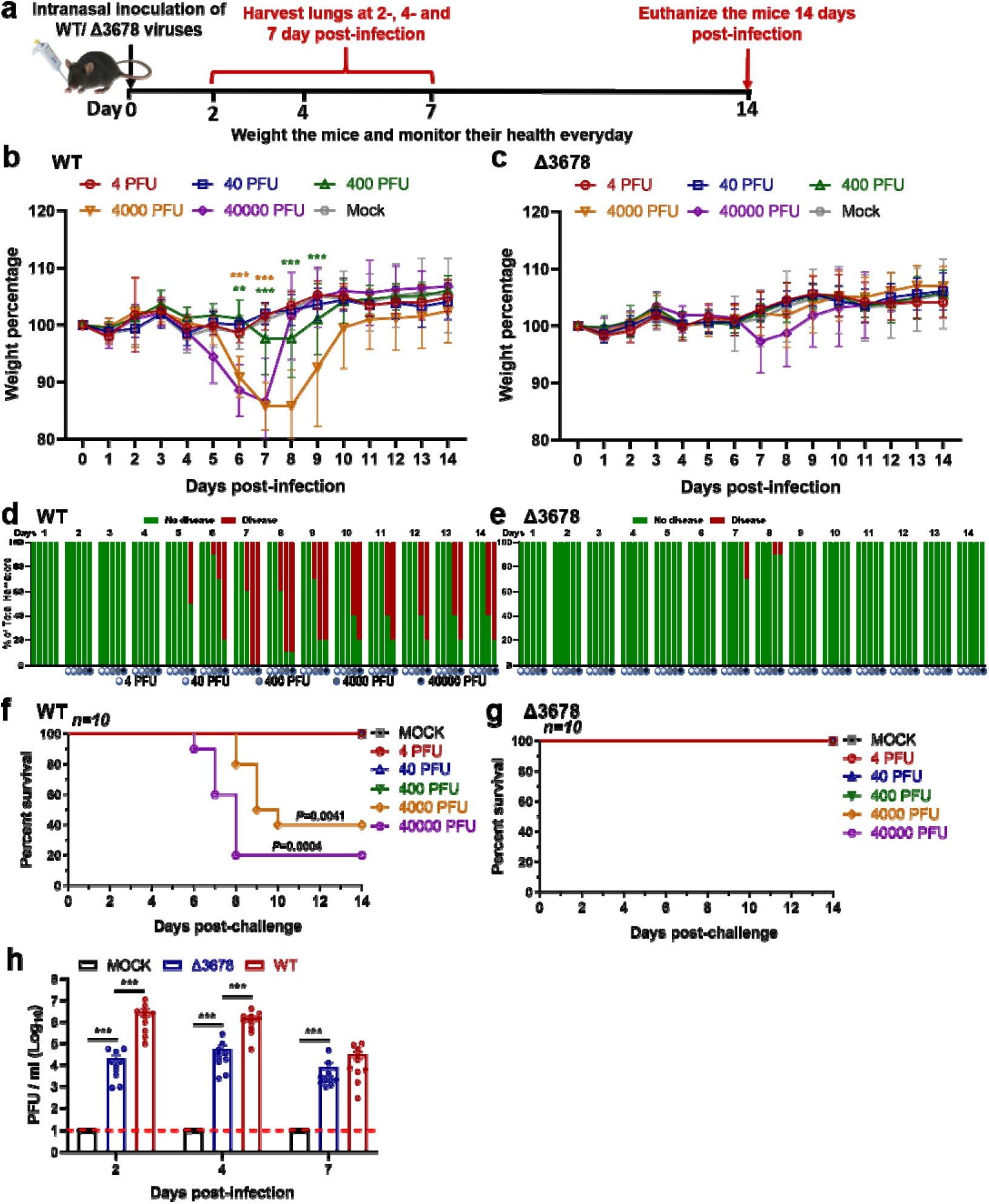
Attenuation of Δ3678 SARS-CoV-2 in K18-hACE2 mice. **a,** Experimental scheme. K-18-hACE2 mice were intranasally inoculated with 4, 40, 400, 4,000, or 40,000 PFU of WT (n=10) or Δ3678 virus (n=10). Lung viral loads were measured on days 2, 4, and 7 post-infection. The infected mice were monitored for body weight (**b,c**), disease (**d,e**), and survivals (**f,g**) for 14 days. The data are shown as mean ± standard deviation. **b,c,** Bodyweight changes. Different viral infection doses are indicated by different colors. The weight changes between mock- and virus-infected groups were statistically analyzed using two-factor analysis of variance (ANOVA) with Tukey’s post hoc test. The green and brown asterisks indicate the statistical difference between mock and 4,000- or 40,000-PFU infection groups. **, P<0.01; ***, P<0.001. **d,e,** Disease. The diseases include ruffled fur, lethargic, hunched posture, or orbital tightening. The percentages of hamsters with or without diseases were presented. **f,g,** Survival. A mixed-model ANOVA using Dunnett’s test for multiple comparisons was used to evaluate the statistical significance. **h,** Lung viral loads from WT- and Δ3678K-infected K18-hACE2 mice. Dots represent individual animals (n = 10). The mean ± standard error of mean is presented. A non-parametric two-tailed Mann-Whitney test was used to determine statistical significance. *P* values were adjusted using the Bonferroni correction to account for multiple comparisons. Differences were considered significant if p<0.025. ***, P<0.0005.

### Genetic stability of Δ3678 SARS-CoV-2 on Vero-E6 cells

Given the potential of Δ3678 as a live-attenuated vaccine, we examined its genetic stability by continuously culturing the virus for five rounds on Vero-E6 cells. Three independent, parallel passaging experiments were performed to assess the consistency of adaptive mutations (**Extended Data Figure 5a**). The passage 5 (P5) virus developed bigger plaques than the original P0 virus (**Extended Data Figure 5b**). Full-genome sequencing of the P5 viruses identified an H655Y substitution in the spike protein and a 675-679 QTQTN spike deletion from all three passage series (**Extended Data Figure 5c**). The substitution and deletion are located immediately upstream of the furin cleavage site between the spike 1 and 2 subunits. Previous studies showed that culturing of SRAS-CoV-2 on Vero cells expressing serine protease TMPRSS2 could eliminate such mutations/deletions^26–28^. Alternatively, to meet the genetic stability required by regulatory agencies, we can engineer the recovered mutations and deletions into our infectious clone to stabilize the vaccine candidate for large-scale production on Vero-E6 cells.

### Contribution of individual ORFs to the attenuation of Δ3678 virus

To define the role of each ORF in attenuating Δ3678 virus, we prepared a panel of mutant viruses in the backbone of a mouse-adapted SARS-CoV-2 (MA-SARS-CoV-2) that can robustly infect BALB/c mice^29^. Each mutant virus contained a single accessory gene deletion, including Δ3a, Δ3b, Δ6, Δ7a, Δ7b, or Δ8. Among all the individual deletion mutants, Δ3a virus developed the smallest plaques on Vero-E6 cells (**Extended Data Figure 6**). The biological importance of each deleted gene was analyzed by viral replication in the lungs after intranasal infection of BALB/c mice (**Fig. 5a**). On day 2 post-infection, deletion of Δ3a, Δ3b, Δ6, Δ7b, or Δ8 reduced viral loads in lungs, among which Δ3a exhibited the largest reduction (**Fig. 5b**). To further confirm the critical role of Δ3a in viral attenuation, we compared the replication kinetics of Δ3a and WT MA-SARS-CoV-2 on Vero-E6 (**Fig. 5c**), Calu-3 (**Fig. 5d**), and HAE cultures (**Fig. 5e**). The replication of Δ3a was significantly more attenuated in immune-competent Calu-3 and HAE cells than in interferon-deficient Vero-E6 cells (**Fig. 5c-e**). Taken together, the results indicate that Δ3a played a major role in attenuating the Δ3678 virus, possibly through the type-I interferon pathway.

**Figure 5.**
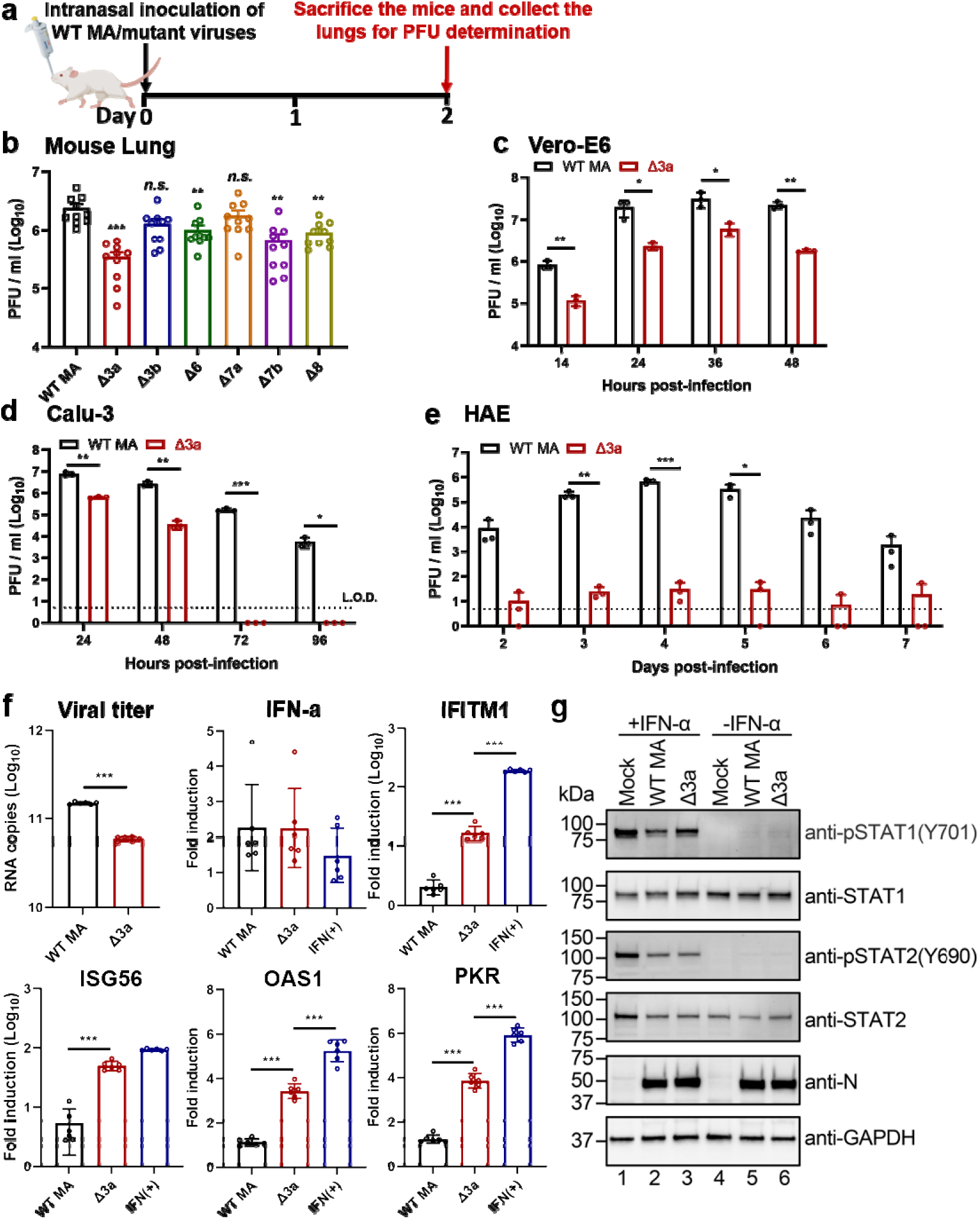
ORF3a deletion is mainly responsible for the attenuation of Δ3678 virus through interfering with STAT1 phosphorylation during type-I interferon signaling. **a,b,** Analysis of individual ORFs in BALB/c mice. **a,** Experimental design. A mouse-adapted SARS-CoV-2 (MA) was used to construct individual ORF-deletion viruses. BALB/c mice were intranasally infected with 10^4^ PFU of WT and individual OFR deletion viruses (n=10 per virus) and quantified for lung viral loads on day 2 post-infection. **b,** Lung viral loads from mice infected with different ORF deletion viruses. Dots represent individual animals (n=10). The mean ± standard error of mean is presented. A non-parametric two-tailed Mann-Whitney test was used to determine the statistical difference between the WT and ORF deletion groups. *P* values were adjusted using the Bonferroni correction to account for multiple comparisons. Differences were considered significant if p<0.0083. *, P<0.0083; **, P<0.0017; ***, P<0.00017. **c-e,** Replication kinetics of Δ3a virus in Vero-E6 (**c**), Calu-3 (**d**), and HAE (**e**) cells. The WT-MA and Δ3a were inoculated onto Vero-E6, Calu-3 and HAE cells at an MOI of 0.01, 0.1, and 2, respectively. After 2 h of incubation, the cells were washed three times with DPBS, continuously cultured with fresh 2% FBS DMEM, and quantified for infectious viruses in culture fluids at indicated time points. Dots represent individual biological replicates (n=3). The values represent the mean ± standard deviation. An unpaired two-tailed *t* test was used to determine significant differences *, P<0.05; **, P<0.01; ***, P<0.001. **f,** ORF3a deletion increases ISG expression in Δ3a-infected A549-hACE2 cells. A549-hACE2 cells were infected with WT or Δ3a virus at an MOI of 1 for 1 h, after which the cells were washed twice with DPBS and continuously cultured in fresh medium. Intracellular RNAs were harvested at 24 h post-infection. Viral RNA copies and mRNA levels of IFN-α, IFITM1, ISG56, OAS1, PKR, and GAPDH were measured by RT-qPCR. The housekeeping gene GAPDH was used to normalize the ISG mRNA levels. The mRNA levels are presented as fold induction over mock samples. As a positive control, uninfected cells were treated with 1,000 units/ml IFN-α for 24 h. Dots represent individual biological replicates (n=6). The data represent the mean ± standard deviation. An unpaired two-tailed *t* test was used to determine significant differences between Δ3a and WT-MA or IFN(+) groups. *P* values were adjusted using the Bonferroni correction to account for multiple comparisons. *, P<0.025; **, P<0.005; ***, P<0.0005. **g,** ORF3a suppresses type-I interferon by reducing STAT1 phosphorylation. A549-hACE2 cells were pre-treated with or without 1,000 units/ml IFN-α for 6 h. The cells were then infected with WT-MA or Δ3a virus at an MOI of 1 for 1 h. The infected cells were washed twice with DPBS, continuously cultured in fresh media with or without 1,000 units/ml IFN-α, and analyzed by Western blot at 24 h post-infection.

### ORF3a antagonizes type-I interferon signaling through inhibiting STAT1 phosphorylation

To define the mechanism of Δ3a-mediated viral attenuation, we infected human lung A549 cells, expressing the human ACE2 receptor (A549-hACE2), with Δ3a or WT MA-SARS-CoV-2. Although the replication of Δ3a was lower than that of the WT virus, comparable levels of IFN-α RNA were produced (**Fig. 5f**). Significantly higher levels of interferon-stimulating genes (ISGs), such as IFITM1, ISG56, OAS1, and PKR, were detected in the Δ3a virus-infected cells than in the WT virus-infected cells (**Fig. 5f**), suggesting a role of ORF3a in suppressing type-I interferon signaling. To further support this conclusion, we treated the A549-hACE cells with IFN-α followed by Δ3a or WT virus infection. Western blot analysis showed that the phosphorylation of STAT1 was less efficient in the Δ3a-infected cells than the WT-infected cells, whereas no difference in STAT2 phosphorylation was observed (**Fig. 5g**). Thus, the results indicate that ORF3a protein suppresses STAT1 phosphorylation during type-I interferon signaling.

### An mNG reporter Δ3678 virus for neutralization and antiviral testing

The *in vitro* and *in vivo* attenuation results suggest that Δ3678 virus may serve as a research tool for BSL-2 use. To further develop this tool, we engineered an mNG gene (driven by its own TRS sequence) between the M and N genes of the Δ3678 genome, resulting in mNG Δ3678 virus (**Fig. 6a**). For high-throughput neutralization testing, we developed the mNG Δ3678 virus into a fluorescent focus reduction neutralization test (FFRNT) in a 96-well format. When infecting Vero-E6 cells, the mNG Δ3678 developed fluorescent foci that could be quantified by high content imaging (**Fig. 6b**). **Fig. 6c** shows the FFRNT curves for one COVID-19 convalescent positive serum one negative serum. To validate the FFRNT assay, we tested 20 convalescent sera against the mNG Δ3678 virus. For comparison, the same serum panel was tested against the WT SARS-CoV-2 (without mNG) using the gold-standard plaque-reduction neutralization test (PRNT)^23^. The 50% reduction neutralization titers (NT_50_) corelated well between the FFRNT and PRNT assays (**Fig. 6d**). The geometric mean of FFRNT_50_/PRNT_50_ ratio was 0.57 for the tested serum panel (**Fig. 6e**). Next, we developed the reporter mNG Δ3678 virus into a high-throughput antiviral assay. Treatment of the mNG Δ3678 virus-infected A549- hACE2 cells with remdesivir diminished the appearance of mNG-positive cells (**Fig. 6f**). Dose-responsive antiviral curves were reliably obtained for the small molecule remdesivir (**Fig. 6g**) and for a monoclonal antibody (**Fig. 6h**). Overall, the results demonstrate that mNG Δ3678 virus could be used for high-throughput neutralization and antiviral testing.

**Figure 6.**
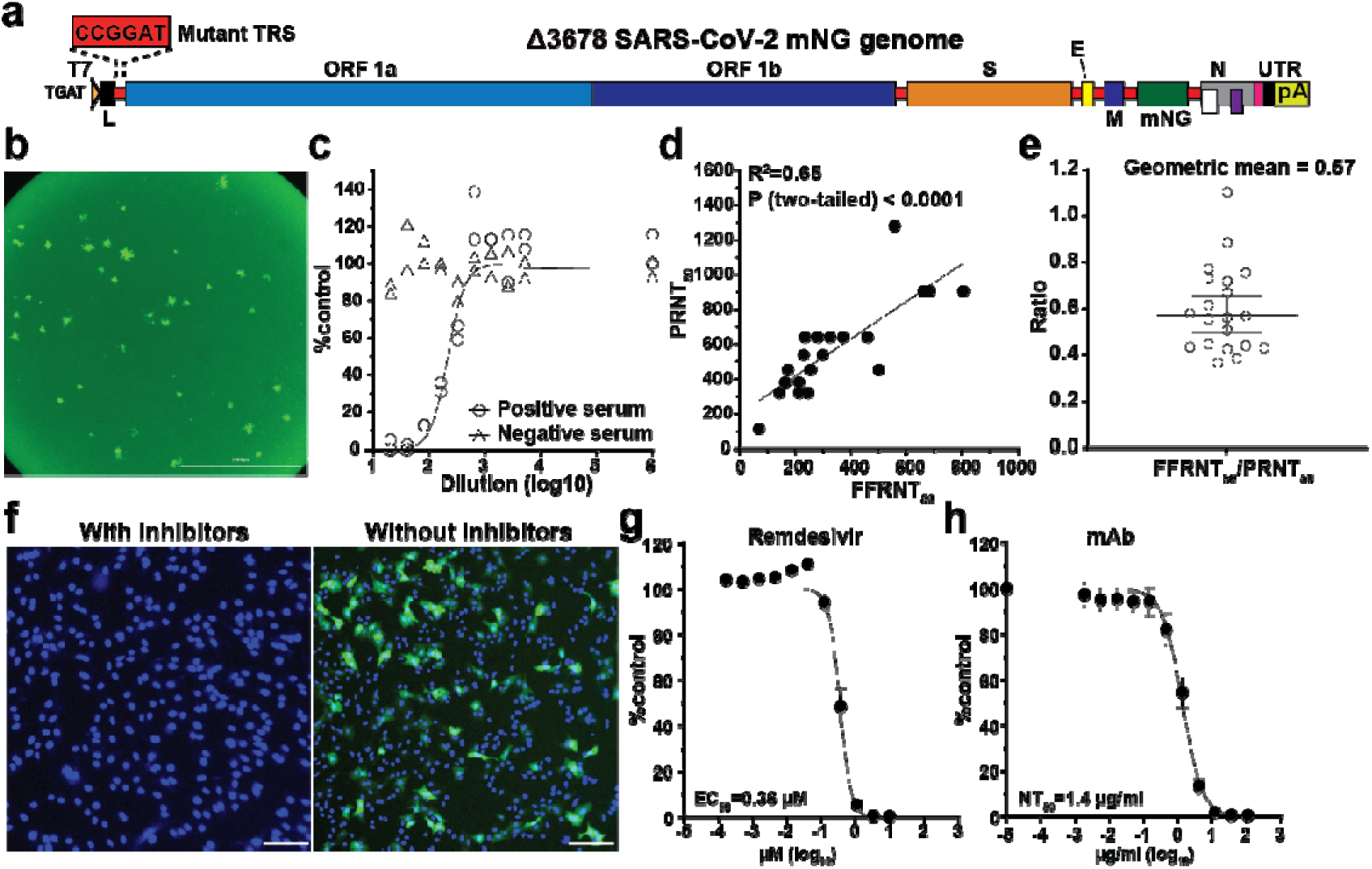
Development of mNG Δ3678 virus for high-throughput neutralization and antiviral testing. **a,** Genome structure of mNG Δ3678 SARS-CoV-2. The mNG gene was inserted between M and N genes. **b**, mNG foci of Vero-E6-TMPRSS2 cells that were infected with mNG Δ3678 SARS-CoV-2 for 16 h. **c,** Representative FFRNT neutralization curves for a COVID-19 antibody-positive and -negative serum. **d**, Correlation between FFRNT_50_ with PRNT_50_ values of 20 COVID-19 convalescent sera. The Pearson correlation efficiency and *P* value are shown. **e**, Scatter-plot of FFRNT_50_/PRNT_50_ ratios. The geometric mean is shown. Error bar indicates the 95% confidence interval of the geometric mean. **f,** Inhibition of mNG-positive cells by remdesivir. A549-hACE2 cells were infected with mNG Δ3678 SARS-CoV-2 in the presence and absence of 10 µM remdesivir. The antiviral activity was measured at 24 h post-infection. **g,** Antiviral response curve of remdesivir against mNG Δ3678 SARS-CoV-2. The calculated 50% effective concentration (EC_50_) is indicated. Error bars indicate the standard deviations from four technical replicates. **h**, The Dose response curve of a monoclonal antibody IgG14. A549-hACE2 cells were infected with Δ3a virus in the presence of different concentrations of IgG14. The mNG signals at 24 h post-infection were used to calculate the NT_50_.

## Discussion

We have developed Δ3678 SARS-CoV-2 as a potential live-attenuated vaccine candidate. The Δ3678 virus could replicate to titers >5.6×10^6^ PFU/ml on interferon-incompetent Vero-E6 cells (**Fig. 1c**), making large-scale production feasible in this vaccine manufacture-approved cell line. In contrast, the Δ3678 virus was highly attenuated when infecting immune-competent cells, as evidenced by the 7,500-fold reduction in viral replication than WT virus on human primary HAE cells (**Fig. 1e**). In both hamster and K18-hACE2 mouse models, the Δ3678 infection did not cause significant weight loss or death at the highest tested infection dose [10^6^ PFU for hamsters (**Fig. 2b**) and 4×10^4^ PFU for K18-hACE2 mice (**Fig. 4c, g**)], whereas the WT virus caused weight loss and death at a much lower infection dose (>4×10^2^ PFU for K18-hACE2 mice; **Fig. 4b, f**). Analysis of individual ORF-deletion viruses identified ORF3a as a major accessory protein responsible for the attenuation of the Δ3678 virus (**Fig. 5b**); this conclusion was further supported by the observation that the addition of Δ3a to the Δ678 virus significantly increased the attenuation of Δ3678 replication (**Fig. 1c-g**). Our results are supported by a recent study reporting that Δ3a SARS-CoV-2 and, to a less extend, Δ6 SARS-CoV-2 were attenuated in the K18 human ACE2 transgenic mice^30^. Mechanistically, we found that ORF3a protein antagonized the innate immune response by blocking STAT1 phosphorylation during type-I interferon signaling. Thus, the deletion of ORF3a conferred SARS-CoV-2 more susceptible to type-I interferon suppression. These findings have uncovered a previously uncharacterized role of ORF3a in the context of SARS-CoV-2 infection. The ORF3a protein was recently shown to form a dimer in cell membrane with an ion channel activity^31^, which may arrcount for its role in cell membrane rearrangement, inflammasome activation, and apoptosis^32–34^. Whether the ion channel activity of ORF3a is required for the inhibition of STAT1 phosphorylation remains to be determined.

The attenuated Δ3678 virus may be pivoted for a veterinarian vaccine. SARS-CoV-2 can infect a variety of animal species, among which cats, ferrets, fruit bats, hamsters, minks, raccoon dogs, and white-tailed deer were reported to spread the infection to other animals of the same species^35–39^, and potentially spillback to humans. A live-attenuated Δ3678 vaccine may be useful for the prevention and control of SARS-CoV-2 on mink farms^40^. Since zoonotic coronaviruses may recombine with the live-attenuated vaccine in immunized animals, we engineered the Δ3678 viruses with mutated TRS to eliminate the possibility of recombination-mediated emergence of WT or replicative chimeric coronaviruses (**Fig. 1a**). This mutated TRS approach was previously shown to attenuate SARS-CoV and to prevent reversion of the WT virus^21, 22^. Given the continuous emergence of SARS-CoV-2 variants, we could update the vaccine antigen by swapping the variant spike glycoproteins into the current Δ3678 virus backbone.

The attenuated Δ3678 virus could serve as a research tool that might be used at BSL-2. Using mNG as an example, we developed an mNG Δ3678 virus for high throughput testing of antibody neutralization and antiviral inhibitors (**Fig. 6**). Depending on research needs, other reporter genes, such as luciferase or other fluorescent genes, could be engineered into the system. This high-throughput assay can be modified for testing neutralization against different variants by swapping the variant spike genes into the Δ3678 backbone. The approach has been successfully used to study vaccine-elicited neutralization against variants in the context of complete SARS-CoV-2^41–43^. Finally, our *in vitro* and *in vivo* attenuation results support the possible use of the Δ3678 virus at BSL-2. If further attenuation is needed, more mutations, such as inactivating the NSP16 2’-O methyltransferase activity^44^, can be rationally engineered into the Δ3678 virus.

One limitation of the current study is that we have not defined the attenuation mechanisms of the ORF 3b, 6, 7b, or 8 deletion, even though they reduced the lung viral loads in the K18- hACE2 mice (**Fig. 5b**). SARS-CoV-2 ORF8 protein was recently reported to contain a histone mimic that could disrupt chromatin regulation and enhance viral replication^45^. Truncations or deletions of ORF7b and ORF8 were reported in SARS-CoV-2 clinical isolates^46, 47^. Future studies are needed to understand the molecular functions of OFR 3b, 6, and 7b proteins. Nevertheless, our results indicate that Δ3678 virus could serve as a live-attenuated vaccine candidate and as an experimental system that can likely be performed at BSL-2 for COVID-19 research and countermeasure development.

## Method

### Ethics statement

Hamster and mouse studies were performed in accordance with the guidance for the Care and Use of Laboratory Animals of the University of Texas Medical Branch (UTMB). The protocol was approved by the Institutional Animal Care and Use Committee (IACUC) at UTMB. All the animal operations were performed under anesthesia by isoflurane to minimize animal suffering. The use of human COVID-19 sera was reviewed and approved by the UTMB Institutional Review Board (IRB#: 20-0070). The convalescent sera from COVID-19 patients (confirmed by the molecular tests with FDA’s Emergency Use Authorization) were leftover specimens and completely de-identified from patient information. The serum specimens were heat-inactivated at 56°C for 30 min before testing.

### Animals and Cells

The Syrian golden hamsters (HsdHan:AURA strain) were purchased from Envigo (Indianapolis, IN). K18-hACE2 mice were purchased from the Jackson Laboratory (Bar Harbor, ME). BALB/c mice were purchased from Charles River Laboratories (Wilmington, MA). African green monkey kidney epithelial Vero-E6 cells (laboratory-passaged derivatives from ATCC CRL-1586) were grown in Dulbecco’s modified Eagle’s medium (DMEM; Gibco/Thermo Fisher, Waltham, MA, USA) with 10% fetal bovine serum (FBS; HyClone Laboratories, South Logan, UT) and 1% antibiotic/streptomycin (P/S, Gibco). Vero-E6-TMPRSS2 cells were purchased from SEKISUI XenoTech, LLC (Kansas City, KS) and maintained in 10% fetal bovine serum (FBS; HyClone Laboratories, South Logan, UT) and 1% P/S and 1 mg/ml G418 (Gibco). The A549-hACE2 cells that stably express hACE2 were grown in the DMEM supplemented with 10% fetal bovine serum, 1% P/S and 1% 4-(2-hydroxyethyl)-1- piperazineethanesulfonic acid (HEPES); ThermoFisher Scientific) and 10 μg/mL Blasticidin S. Human lung adenocarcinoma epithelial Calu-3 cells (ATCC, Manassas, VA, USA) were maintained in a high-glucose DMEM containing sodium pyruvate and GlutaMAX (Gibco) with 10% FBS and 1% penicillin/streptomycin at 37°C with 5% CO_2_. The EpiAirway system is a primary human airway 3D tissue model purchased from MatTek Life Science (Ashland, MA, USA). All cells were maintained at 37°C with 5% CO_2_. All cell lines were verified and tested negative for mycoplasma.

### Generation of SARS-CoV-2 mutant viruses

(1) Generate mutant viruses with accessory ORF deletions. The ORF 6, 7, and 8 deletions (Δ678) and ORF 3, 6, 7, and 8 deletions (Δ3678) were constructed by overlap PCR using an infection clone of USA-WA1/2020 SARS-CoV-2^8^. The Δ3a, Δ3b, Δ6, Δ7a, Δ7b, Δ8 mutants were engineered into an infection clone of a mouse-adapted SARS-CoV-2 (MA-SARS-CoV-2)^29^ using a standard molecular cloning protocol. For generating ΔORF6 and ΔORF8 mutants, an overlapping PCR strategy was used to delete the ORF and the upstream transcriptional regulatory sequence (TRS). For generating ΔORF3a and ΔORF7a mutants, the complete ORF3 and ORF7 were replaced with the ORF3b- and ORF7b-coding sequence, respectively. For generating ΔORF3b mutant, several nonsense mutations were introduced into MA-SARS-CoV-2 to disrupt the initiation codon of ORF3b without affecting the translation of ORF3a (*i.e.*, the change of sequence from wild-type *T<u>A**T**GA**T**G</u>* to mutant *T<u>A**C**GA**C**G</u>*; ORF3b- coding sequence is underlined). For producing ΔORF7b mutant, the initiation codon of ORF7b was disrupted, and the ORF7b gene was deleted from the fifth nucleotide position onward. (2) Generate reporter viruses with accessory ORF deletions. The mNG WT and mNG Δ3678 SARS-CoV-2s were generated by engineering the mNeonGreen (mNG) gene into the ORF7 position of the WT and Δ3678 viruses. The mutant infectious clones were assembled by *in vitro* ligation of contiguous DNA fragments following the protocol previously described^48^. *In vitro* transcription was then performed to synthesize genomic RNA. For recovering the viruses, the RNA transcripts were electroporated into Vero-E6 cells. The viruses from electroporated cells were harvested at 40 h post-electroporation and served as P0 stocks. All viruses were passaged once on Vero-E6 cells for subsequent experiments and sequenced after RNA extraction to confirm no undesired mutations. Viral titers were determined by plaque assay on Vero-E6 cells. All virus preparation and experiments were performed in a BSL-3 facility. Viruses and plasmids are available from the World Reference Center for Emerging Viruses and Arboviruses (WRCEVA) at the University of Texas Medical Branch.

### RNA extraction, RT-PCR, and cDNA sequencing

Cell culture supernatants or clarified tissue homogenates were mixed with a five-fold excess of TRIzol™ LS Reagent (Thermo Fisher Scientific, Waltham, MA). Viral RNAs were extracted according to the manufacturer’s instructions. The extracted RNAs were dissolved in 20 μl nuclease-free water. For sequence validation of mutant viruses, 2 µl of RNA samples were used for reverse transcription by using the SuperScript™ IV First- Strand Synthesis System (Thermo Fisher Scientific) with random hexamer primers. Nine DNA fragments flanking the entire viral genome were amplified by PCR. The resulting DNAs were cleaned up by the QIAquick PCR Purification Kit, and the genome sequences were determined by Sanger sequencing at GENEWIZ (South Plainfield, NJ).

### Viral infection of cell lines

Approximately 3×10^5^ Vero-E6 or Calu-3 cells were seeded onto each well of 12-well plates and cultured at 37°C, 5% CO_2_ for 16 h. SARS-CoV-2 WT or mutant viruses were inoculated onto Vero-E6 and Calu-3 cells at an MOI of 0.01 and 0.1, respectively. After 2 h infection at 37°C with 5% CO_2_, the cells were washed with DPBS 3 times to remove any detached virus. One milliliter of culture medium was added to each well for the maintenance of the cells. At each time point, 100 µl of culture supernatants were collected for detection of virus titer, and 100 µl of fresh medium was added into each well to replenish the culture volume. The cells were infected in triplicate for each group of viruses. All samples were stored at -80°C until analysis.

### Viral infection in a primary human airway cell culture model

The EpiAirway system is a primary human airway 3D mucociliary tissue model consisting of normal, human-derived tracheal/bronchial epithelial cells. For viral replication kinetics, WT or mutant viruses were inoculated onto the culture at an indicated MOI in DPBS. After 2 h infection at 37°C with 5% CO_2_, the inoculum was removed, and the culture was washed three times with DPBS. The infected epithelial cells were maintained without any medium in the apical well, and the medium was provided to the culture through the basal well. The infected cells were incubated at 37°C, 5% CO_2_. From 1-7 days, 300 μl of DPBS were added onto the apical side of the airway culture and incubated at 37°C for 30 min to elute the released viruses. All virus samples in DPBS were stored at −80°C.

### Quantitative real-time RT-PCR assay

RNA copies of SARS-CoV-2 samples were detected by quantitative real-time RT-PCR (RT-qPCR) assays were performed using the iTaq SYBR Green One-Step Kit (Bio-Rad) on the LightCycler 480 system (Roche, Indianapolis, IN) following the manufacturer’s protocols. The absolute quantification of viral RNA was determined by a standard curve method using an RNA standard (*in vitro* transcribed 3,480 bp containing genomic nucleotide positions 26,044 to 29,883 of SARS-CoV-2 genome).

### Hamster immunization and challenge assay

Four- to six-week-old male golden Syrian hamsters, strain HsdHan:AURA (Envigo, Indianapolis, IN), were immunized intranasally with 100 µl WT virus (10^6^ PFU, n=20) or Δ3678 mutant virus (10^6^ PFU, n=20). Animals received DMEM media (supplemented with 2% FBS and 1% penicillin/streptomycin) served as Mock group (n=20). On day 28, the animals were challenged with 10^5^ PFU of WT SARS-CoV-2. The animals were weighed and monitored for signs of illness daily. Nasal washes and oral swabs were collected in 400 µl sterile DPBS and 1 ml DMEM media at indicated time points. For organ collection, animals were humanely euthanized on days 2, 30, and 32, tracheae and lungs were harvested and placed in a 2-ml homogenizer tube containing 1 ml of DMEM media. On day 49, animals were humanely euthanized for blood collection, serum were then isolated for neutralization titer (NT_50_) detection. The NT_50_ values were determined using an mNG USA-WA1/2020 SARS-CoV-2 as previously reported^23^.

To test if lower dose immunization could achieve protection, hamsters were immunized with 10^2^, 10^3^, 10^4^, or 10^5^ PFU of Δ3678 virus. Nasal washes, oral swabs and organs were collected at indicated time points. The animals were weighed and monitored for signs of illness daily.

### Hamster transmission assay

Hamster transmission assay was performed per our previous protocol^24^. Briefly, hamsters were immunized intranasally with 10^6^ PFU Δ3678 mutant virus (n=5). Animals who received DMEM media served as a mock group (n=5). On day 28 post-immunization, the animals were challenged with 10^5^ PFU of WT SARS-CoV-2. One day later, one infected donor animal was co-housed with one naïve animal for 8 h (5 pairs for mock group, 5 pairs for Δ3678 group). After the 8-h contact, the donor and recipient animals were separated and maintained in individual cages. Nasal washes were collected at indicated time points. On day 42, animals were humanely euthanized for blood collection.

### Mouse infection of Δ3678 virus

Eight- to 10-week-old K18-hACE2 female mice were intranasally infected with 50 µl different doses of WT or Δ3678 virus (4, 40, 400, 4,000, 40,000 PFU, n=10 per dose). Animals received DMEM media served as a mock group. Lungs were collected on days 2, 4, and 7 post-infection. Animals were weighed and monitored for signs of illness daily and were sacrificed on day 14.

To define the role of each ORF in attenuating Δ3678 virus, 8- to 10-week-old BALB/c female mice were intranasally infected with 50 µl WT mouse-adapted-SARS-CoV-2 (10^4^ PFU, n=10) or Δ3a, Δ3b, Δ6, Δ7a, Δ7b, Δ8 virus (10^4^ PFU, n=10 per virus). On day 2 post-infection, animals were humanely euthanized for lung collection.

### Histopathology

Hamsters were euthanized with ketamine/xylazine injection and necropsy was performed. The lungs were inspected for gross lesions and representative portions of the lungs were collected in 10% buffered formalin for histology. Formalin-fixed tissues were processed per a standard protocol, 4 μm-thick sections were cut and stained with hematoxylin and eosin (HE). The slides were imaged in a digital scanner (Leica Aperio LV1). Lung sections were examined under light microscopy using an Olympus CX43 microscope for the extent of inflammation, size of inflammatory foci, and changes in alveoli, alveolar septa, airways, and blood vessels. The blinded tissue sections were semi-quantitatively scored for pathological lesions.

### Plaque assay

Approximately 1.2×10^6^ Vero-E6 cells were seeded to each well of 6-well plates and cultured at 37°C, 5% CO_2_ for 16 h. Virus was serially diluted in DMEM with 2% FBS and 200 µl diluted viruses were transferred onto the monolayers. The viruses were incubated with the cells at 37°C with 5% CO_2_ for 1 h. After the incubation, overlay medium was added to the infected cells per well. The overlay medium contained DMEM with 2% FBS, 1% penicillin/streptomycin, and 1% sea-plaque agarose (Lonza, Walkersville, MD). After a 2-day incubation, the plates were stained with neutral red (Sigma-Aldrich, St. Louis, MO) and plaques were counted on a lightbox. The detection limit of the plaque assay was 10 PFU/ml.

### Genetic stability of Δ3678 SARS-CoV-2

The P0 Δ3678 SARS-CoV-2 was continuously cultured for five rounds on Vero-E6 cells. Three independent passaging experiments were performed to assess the consistency of adaptive mutations. The P5 viral RNAs from three independent replicates were extracted and subjected to RT-PCR. Whole-genome sequencing was performed on RT-PCR products. The mutations that occurred in the P5 Δ3678 viruses were analyzed.

### ORF3a-mediated suppression of type-I interferon signaling

A549-hACE2 cells were infected with WT or ΔORF3a SARS-CoV-2 at an MOI of 1 for 1 h, after which the cells were washed twice with PBS and cultured in a fresh medium. Intracellular RNAs were harvested at 24 h post-infection. Viral RNA copies and mRNA levels of IFN-α, IFITM1, ISG56, OAS1, PKR, and GAPDH were determined by quantitative RT-PCR. The housekeeping gene GAPDH was used to normalize mRNA levels and the mRNA levels are presented as fold induction over mock samples. As a positive control, uninfected cells were treated with 1,000 units/ml IFN-α for 24 h.

### Suppression of STAT1 phosphorylation by ORF3a protein

A549-hACE2 cells were pre-treated with 1,000 units/ml IFN-α for 6 h. Mock-treated cells were used as a control. Cells were infected with WT or ΔORF3 SARS-CoV-2 at an MOI 1 for 1 h. Inoculums were removed; cells were washed twice with PBS; fresh media with or without 1,000 units/ml IFN-α were added. Samples were collected at 24 h post-infection by using 2 x Laemmli buffer (BioRad, #1610737) and analyzed by Western blot. Recombinant human α-interferon (IF007) was purchased from Millipore (Darmstadt, Germany). Anti-STAT1 (14994S, 1:1,000), anti-pSTAT1 (Y701) (7649S, 1:1,000), anti-STAT2 (72604S, 1:1,000), anti-pSTAT2 (Y690) (88410S, 1:1,000) antibodies were from Cell Signaling Technology (Danvers, MA); anti-GAPDH (G9545, 1:1,000) antibodies were from Sigma-Aldrich; SARS-CoV-2 (COVID-19) nucleocapsid antibody (NB100-56576, 1:1000) were from Novus Biologicals (CO, USA).

### Fluorescent focus reduction neutralization test (FFRNT)

Neutralization titers of COVID-19 convalescent sera were measured by a fluorescent focus reduction neutralization test (FFRNT) using mNG Δ3678 SARS-CoV-2. Briefly, Vero-E6 cells (2.5 × 10^4^) were seeded in each well of black μCLEAR flat-bottom 96-well plate (Greiner Bio-one™). The cells were incubated overnight at 37°C with 5% CO2. On the following day, each serum was 2-fold serially diluted in the culture medium with the first dilution of 1:20. Each serum was tested in duplicates. The diluted serum was incubated with 100-150 fluorescent focus units (FFU) of mNG SARS-CoV-2 at 37°C for 1 h (final dilution range of 1:20 to 1:20,480), after which the serum-virus mixtures were inoculated onto the pre-seeded Vero-E6 cell monolayer in 96-well plates. After 1 h infection, the inoculum was removed and 100 μl of overlay medium (DMEM supplemented with 0.8% methylcellulose, 2% FBS, and 1% P/S) was added to each well. After incubating the plates at 37°C for 16 h, raw images of mNG fluorescent foci were acquired using CytationTM 7 (BioTek) armed with 2.5× FL Zeiss objective with widefield of view and processed using the software settings (GFP [469,525] threshold 4000, object selection size 50-1000 µm). The foci in each well were counted and normalized to the non-serum-treated controls to calculate the relative infectivities. The curves of the relative infectivity versus the serum dilutions (log10 values) were plotted using Prism 9 (GraphPad). A nonlinear regression method with log (inhibitor) vs. response-variable slope (four parameters) model (bottom and top parameters were constrained to 0 and 100, respectively) was used to determine the dilution fold that neutralized 50% of mNG SARS-CoV-2 (defined as FFRNT_50_) in GraphPad Prism 9. Each serum was tested in duplicates.

### Antiviral testing

A549-hACE2 cells were used to evaluate the efficacy of a monoclonal antibody IgG14 and antiviral drug remdesivir. The sources of IgG14 and remdesivir were previously reported^9, 49^. Briefly, A549-hACE2 cells (1.2 × 10^4^) were seeded in each well of black μCLEAR flat-bottom 96-well plate (Greiner Bio-one™). The cells were incubated overnight at 37°C with 5% CO_2_. For antibody testing, on the following day, IgG14 was 3- fold serially diluted and incubated with mNG Δ3678 at 37°C for 1 h, after which the antibody-virus mixtures were inoculated into the 96-well plates that were pre-seeded A549-hACE2 cells. For antiviral testing, remdesivir was 3-fold serially diluted in DMSO and further diluted as 100 folds in the culture medium containing mNG Δ3678 virus. Fifty µl of the compound-virus mixture were immediately added to the cells at a final MOI of 1.0. At 24 h post-infection, 25 μl of Hoechst 33342 Solution (400-fold diluted in Hank’s Balanced Salt Solution; Gibco) were added to each well to stain the cell nucleus. The plate was sealed with Breath-Easy sealing membrane (Diversified Biotech), incubated at 37°C for 20 min, and quantified for mNG fluorescence on CX5 imager (ThermoFisher Scientific). The raw images (2 × 2 montage) were acquired using 4× objective. The total cells (indicated by nucleus staining) and mNG-positive cells were quantified for each well. Infection rates were determined by dividing the mNG-positive cell number by total cell number. Relative infection rates were obtained by normalizing the infection rates of treated groups to those of no-treated controls. The curves of the relative infection rates versus the concentration were plotted using Prism 9 (GraphPad). A nonlinear regression method was used to determine the concentration of antiviral that suppress 50% of mNG fluorescence (EC_50_). Experiment was tested in quadruplicates.

### Statistics

Hamsters and mice were randomly allocated into different groups. The investigators were not blinded to allocation during the experiments or to the outcome assessment. No statistical methods were used to predetermine sample size. Descriptive statistics have been provided in the figure legends. For *in vitro* replication kinetics, Kruskal–Wallis analysis of variance was conducted to detect any significant variation among replicates. If no significant variation was detected, the results were pooled for further comparison. Differences between continuous variables were assessed with a non-parametric Mann– Whitney test. The PFU and genomic copies were analyzed using an unpaired two-tailed *t* test. The weight loss data were shown as mean ± standard deviation and statistically analyzed using two-factor analysis of variance (ANOVA) with Tukey’s post hoc test. The animal survival rates were analyzed using a mixed-model ANOVA using Dunnett’s test for multiple comparisons. Analyses were performed in Prism version 9.0 (GraphPad, San Diego, CA).

## Acknowledgements

We thank the colleagues at University of Texas Medical Branch (UTMB) for helpful discussions during the course of this work. P.-Y.S. was supported by NIH grants HHSN272201600013C, AI134907, AI145617, and UL1TR001439, and awards from the Sealy & Smith Foundation, the Kleberg Foundation, the John S. Dunn Foundation, the Amon G. Carter Foundation, the Gilson Longenbaugh Foundation, and the Summerfield Robert Foundation. V.D.M. was supported by NIH grants U19AI100625, R00AG049092, R24AI120942, and a STARs Award from the University of Texas System. S.C.W. was supported by NIH grant R24 AI120942. A.B. was supported by Defense Advanced Research Project Agency grant N6600119C4022 and NIH grant U19 AI142790. T.W. was supported by NIH grants R01AI127744 and R21 AI140569. J.L. was supported by the postdoctoral fellowship from the McLaughlin Fellowship Endowment at UTMB.

## Author contributions

V.D.M., X.X., and P.-Y.S. conceived the study. Y.L., X.Z., J.L., H.X., J.Z., A.E.M., S.P., J.A.P., N.E.B., C.K., K.S.P., and X.X. performed the experiments. Y.L., X.Z., J.L., H.X., S.P. N.E.P., A.B., P.R., V.D.M., K.S.P., X.X., S.C.W., and P.-Y.S. analyzed the results. V.D.M. and P.R. provided critical reagents. Y.L., X.Z., J.L., H.X., V.D.M., K.S.P., X.X., S.C.W., and P.-Y.S. wrote the manuscript.

## Data Availability

The results presented in the study are available upon request from the corresponding authors. The mNG reporter Δ3678 SARS-CoV-2 will be deposited to the World Reference Center for Emerging Viruses and Arboviruses (https://www.utmb.edu/wrceva) at UTMB for distribution.

---

**Extended Data Figure 1.**
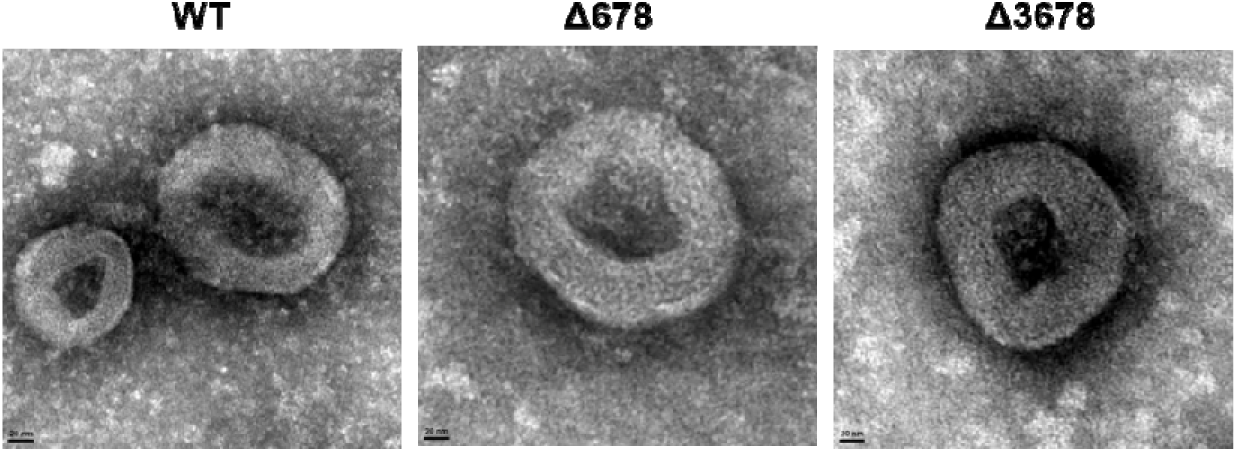
Negative-staining electron microscopic images of WT, Δ678, and Δ3678 SARS-CoV-2s. Scale bar, 20 nm.

**Extended Figure 2.**
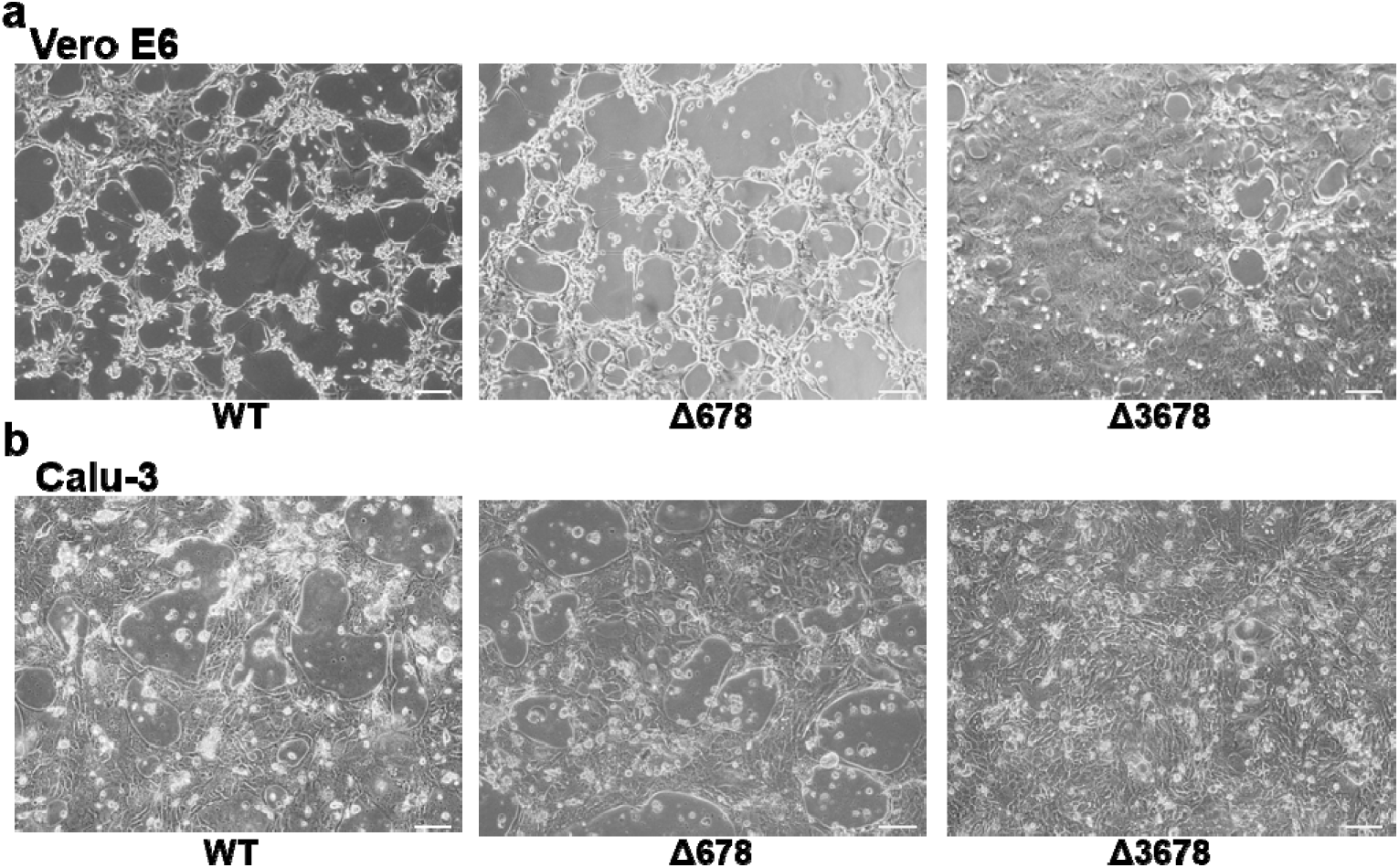
Brightfield images of the cytopathic effects of WT, Δ678, and Δ3678 SARS-CoV-2-infected Vero-E6 and Calu-3 cells. The Vero-E6 and Calu-3 cells were infected with WT, Δ678, or Δ3678 virus at an MOI of 0.1 and 1.0, respectively. The images of infected Vero-E6 (**a**) and Calu-3 cells (**b**) were taken at 24 and 48 h post-infection, respectively. Scale bar, 100 µm.

**Extended Data Figure 3.**
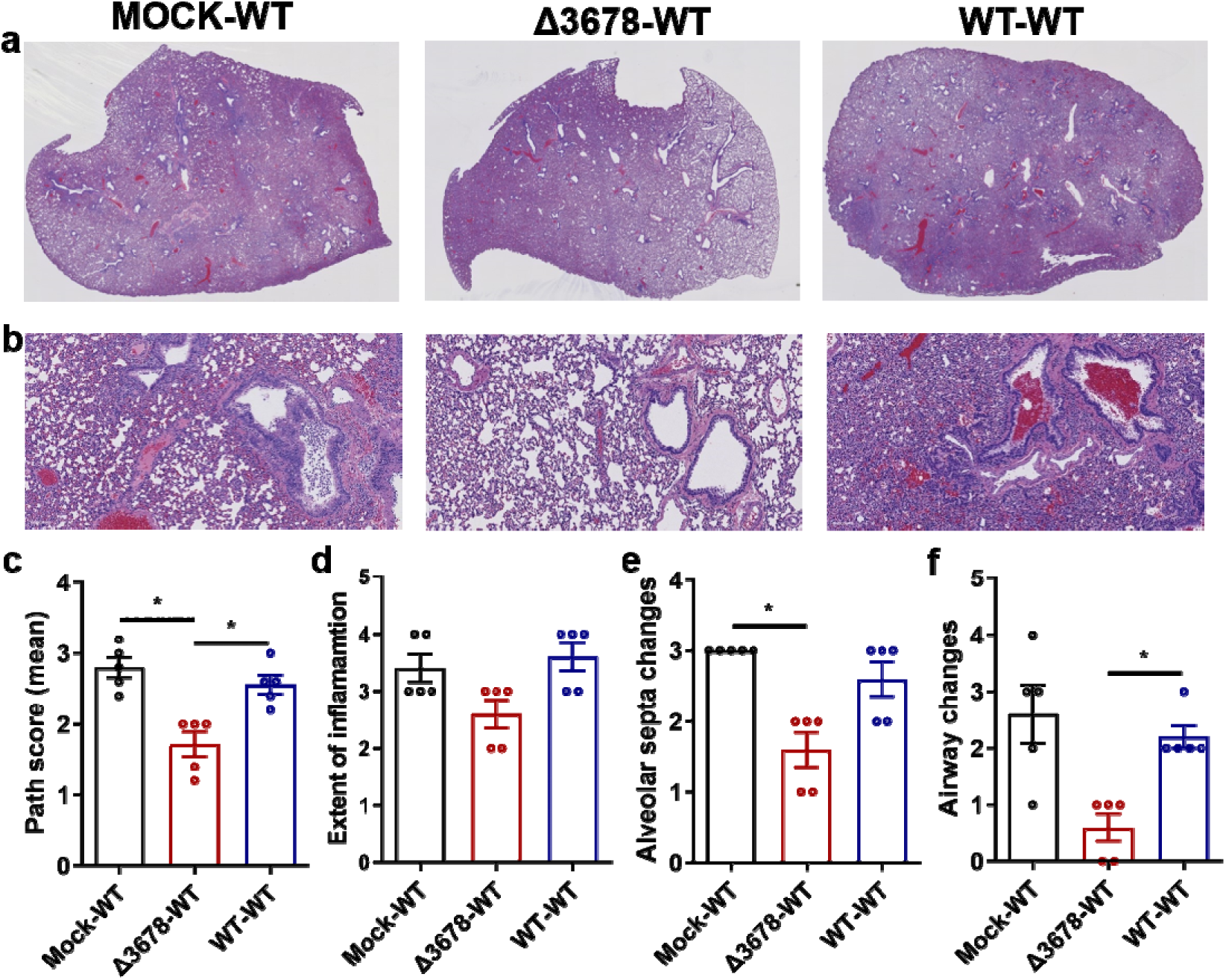
Lung pathology of Δ3678 virus-immunized and WT SARS-CoV-2-challenged hamsters. **a,** Lung sections show typical interstitial pneumonia with moderate to severe inflammatory changes in mock-immunized and WT virus-challenged animals (mock-WT) or in WT virus-inoculated and WT virus-challenged animals (WT-WT). Reduced inflammatory changes are observed in Δ3678 virus-immunized and WT virus-challenged animals (Δ3678-WT). **b,** Higher magnification images show large inflammatory cells in the airways and prominent septal thickening in the mock-WT and WT-WT groups. Such changes are minimal or absent in the Δ3678-WT group. **c-f,** Comparative pathology scores calculated based on the criteria described in Table S1. The Δ3678-WT group shows a significant reduction in total pathology score (**c**), extent of inflammation (**d**), alveolar septa changes (**e**), and airway changes (**f**). Dots represent individual animals (n=5). The values of mean ± standard error of mean are presented. A non-parametric two-tailed Mann-Whitney test was used to determine the statistical differences between Δ3678-WT and mock-WT or WT-WT groups. *P* values were adjusted using the Bonferroni correction to account for multiple comparisons. *, P<0.025.

**Extended Data Figure 4.**
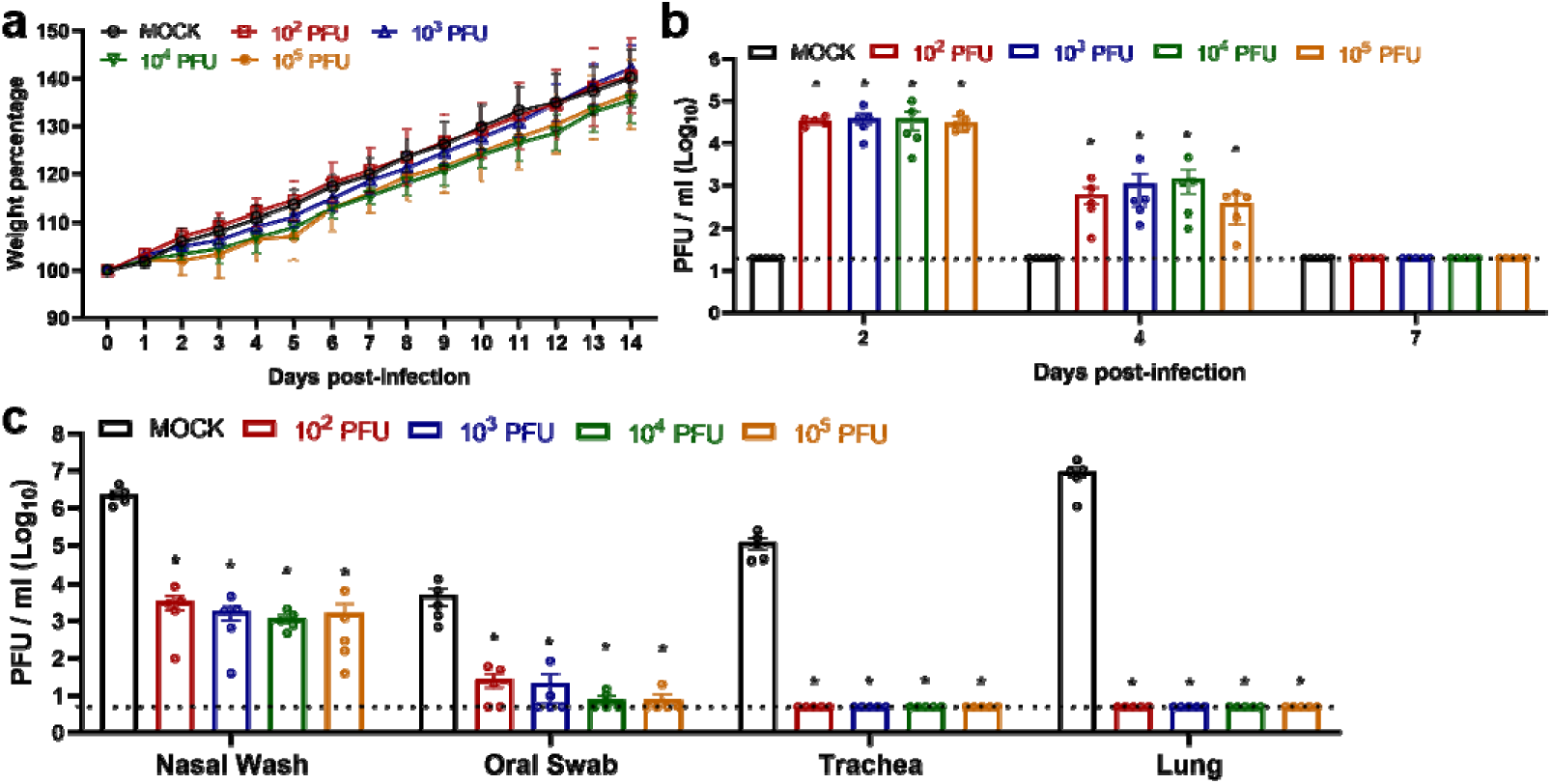
Dose range immunization of Δ3678 virus to protect hamsters from WT SARS-CoV-2 challenge. **a,** Weight loss of hamsters immunized with four different doses of Δ3678 virus. Hamsters were intranasally inoculated with 10^2^, 10^3^, 10^4^, or 10^5^ PFU of Δ3678 virus (n=5 per dose). Body weights were measured for 14 days post-inoculation. The data are shown as mean ± standard deviation. The weight changes were statistically analyzed using two-factor analysis of variance (ANOVA) with Tukey’s post hoc test. No statistic differences were observed among mock and all Δ3678 dose groups. **b,** Nasal viral loads in Δ3678 virus-immunized hamsters on days 2, 4, and 7 post-immunization. **c,** Viral loads in nasal wash, oral swab, trachea, and lung from Δ3678-immunized and WT virus-challenged hamsters. The Δ3678-immunized hamsters were challenged with WT SARS-CoV-2 on day 28 post-immunization. The viral loads were measured on day 2 post-challenge. **b,c,** Dots represent individual animals (n=5). The values in the graph represent the mean ± standard error of mean. Dash lines indicate assay detection limitations. A non-parametric two-tailed Mann-Whitney test was used to determine the statistical differences between mock- and Δ3678-immunized hamsters. *P* values were adjusted using the Bonferroni correction to account for multiple comparisons. Differences were considered significant if p<0.0125. *, P<0.0125.

**Extended Data Figure 5.**
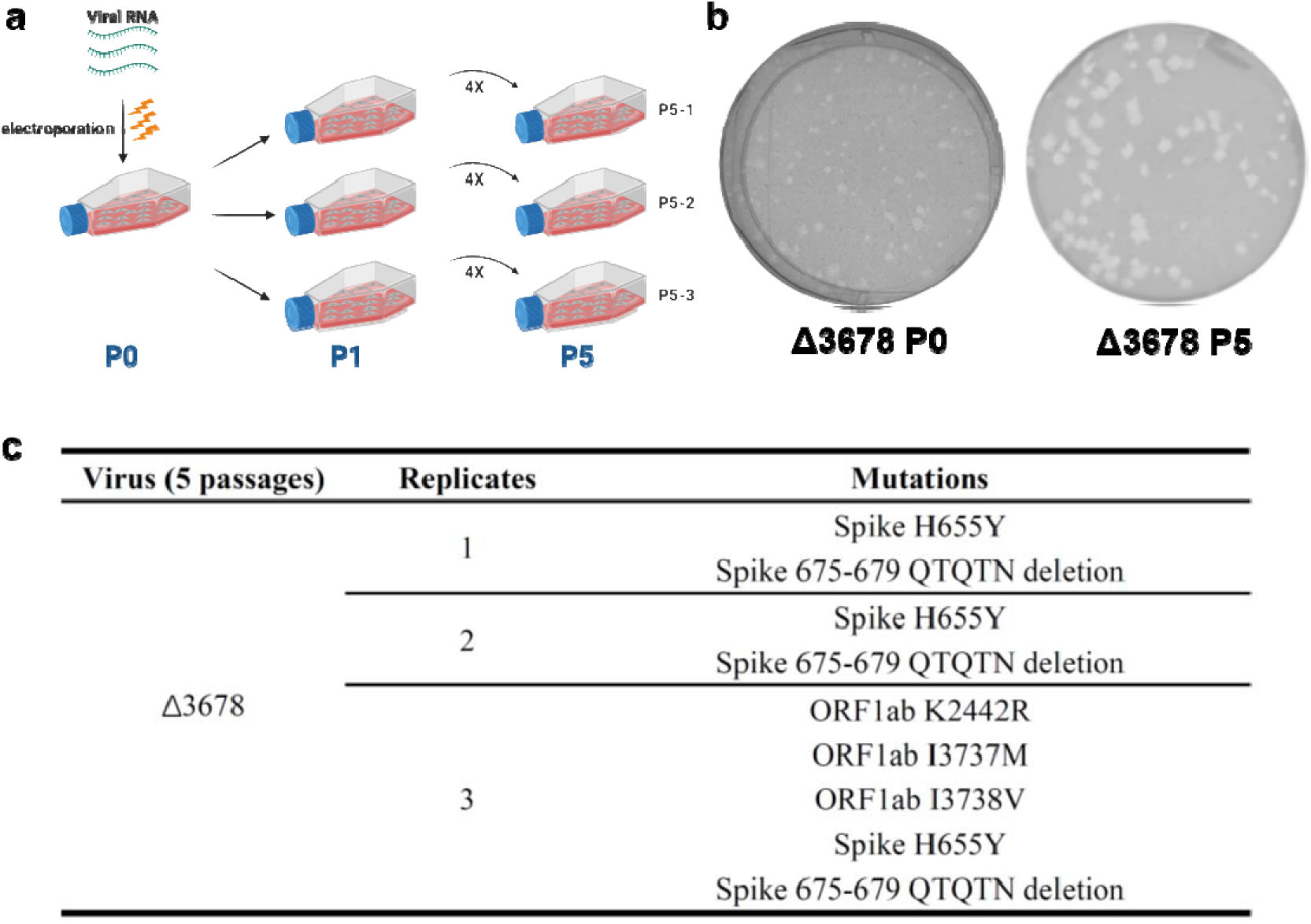
Genetic stability analysis of Δ3678 SARS-CoV-2. **a,** Experimental scheme. Passage 0 (P0) Δ3678 virus was divided into three T25 flasks for five rounds of independent passaging on Vero-E6 cells. **b,** Plaque morphologies of P0 and P5 Δ3678 virus. **c,** Mutations recovered from three independently cultured P5 Δ3678 viruses. The P5 Δ3678 viral RNAs were extracted and amplified by RT-PCR. Whole-genome sequencing was performed on the RT-PCR products. Mutations from the P5 viruses were annotated with amino acid changes and specific genes.

**Extended Data Figure 6.**
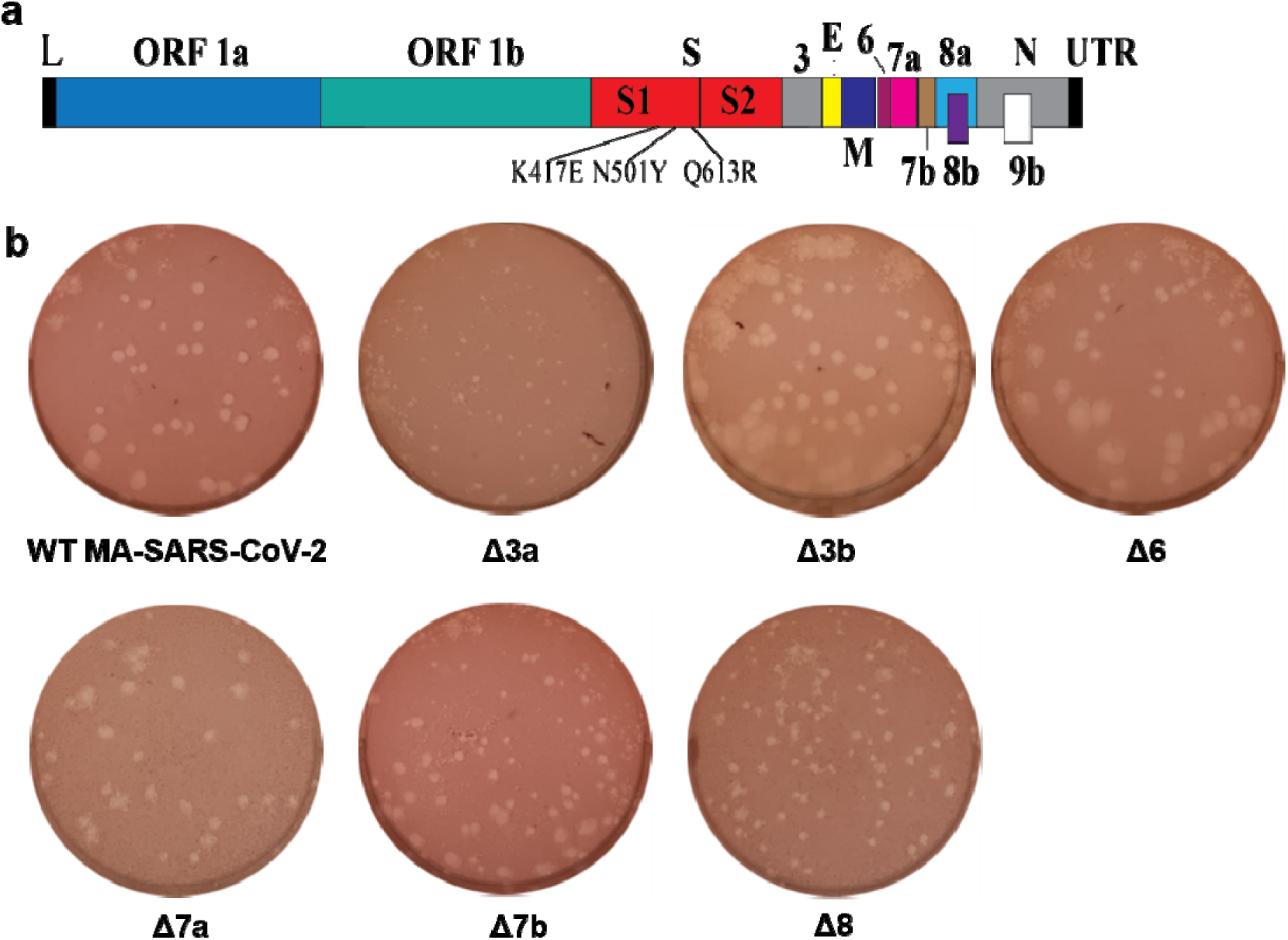
Construction of mouse-adapted SARS-CoV-2s with individual ORF deletions. **a,** Mouse-adapted SARS-CoV-2 genome. Mouse-adapted SARS-CoV-2 (MA-SARS-CoV-2) contains three mutations in spike glycoprotein: K417E, N501Y, and Q613R. These mutations confer SARS-CoV-2 to replicate in BALB/c mice. Open reading frames, ORFs; E, envelope glycoprotein gene; L, leader sequence; M, membrane glycoprotein gene; N, nucleocapsid gene; UTR, untranslated region. **b,** Plaque morphologies of MA-SARS-CoV-2s with individual ORF deletions. All these ORF deletion viruses were constructed in the backbone of MA-SARS-CoV-2. Plaque assays were performed on Vero-E6 cells and stained on day 2.5 post-infection.

**Extended Data Table 1.**
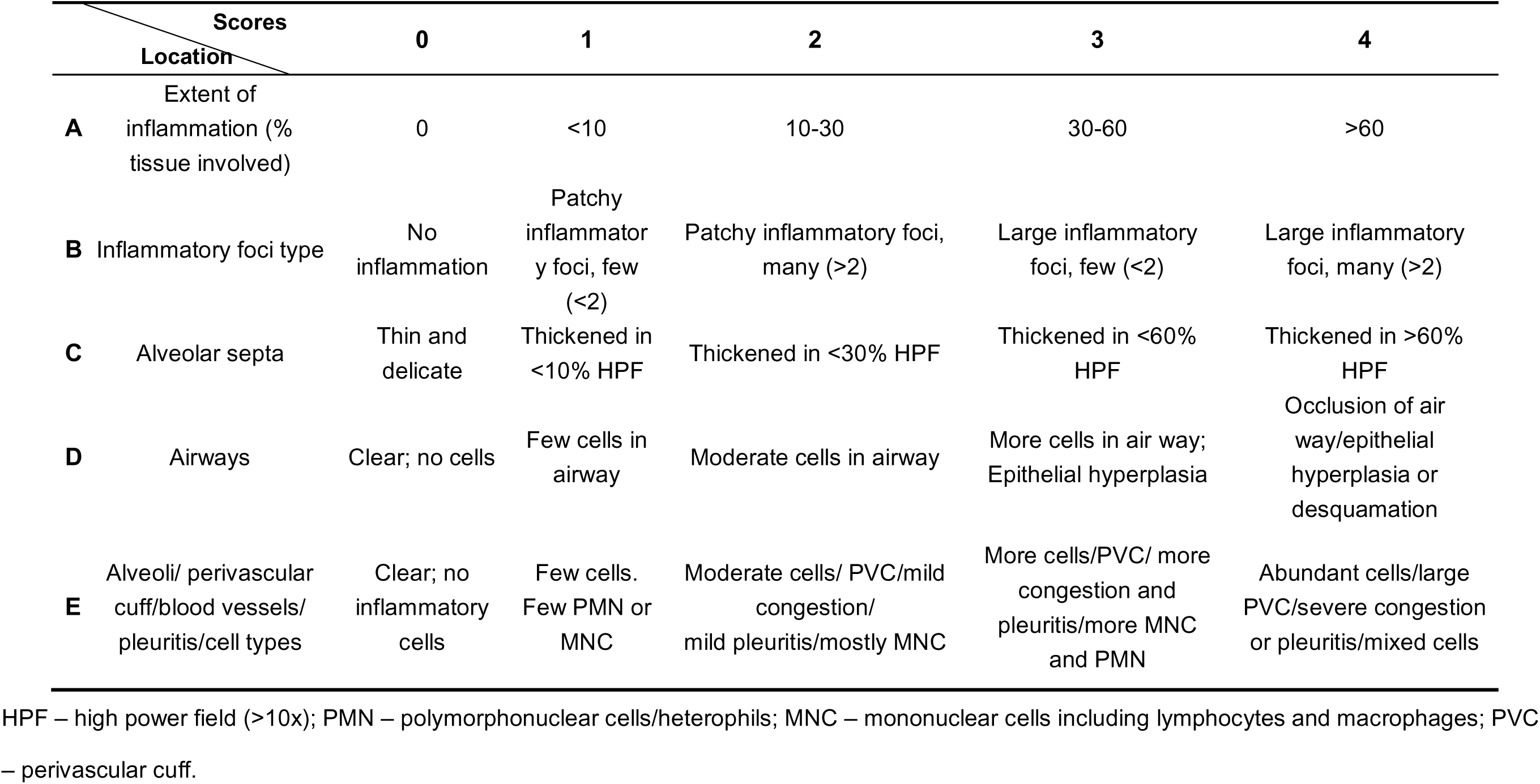
Criteria for histopathology scoring.

| Scores |  | 0 | 1 | 2 | 3 | 4 |
| --- | --- | --- | --- | --- | --- | --- |
| Location |  |  |  |  |  |  |
| **A** | Extent of inflammation (% tissue involved) | 0 | <10 | 10-30 | 30-60 | >60 |
| **B** | Inflammatory foci type | No inflammation | Patchy inflammatory foci, few (<2) | Patchy inflammatory foci, many (>2) | Large inflammatory foci, few (<2) | Large inflammatory foci, many (>2) |
| **C** | Alveolar septa | Thin and delicate | Thickened in <10% HPF | Thickened in <30% HPF | Thickened in <60% HPF | Thickened in >60% HPF |
| **D** | Airways | Clear; no cells | Few cells in airway | Moderate cells in airway | More cells in air way; Epithelial hyperplasia | Occlusion of air way/epithelial hyperplasia or desquamation |
| **E** | Alveoli/ perivascular cuff/blood vessels/ pleuritis/cell types | Clear; no inflammatory cells | Few cells. Few PMN or MNC | Moderate cells/ PVC/mild congestion/ mild pleuritis/mostly MNC | More cells/PVC/ more congestion and pleuritis/more MNC and PMN | Abundant cells/large PVC/severe congestion or pleuritis/mixed cells |

HPF – high power field (>10x); PMN – polymorphonuclear cells/heterophils; MNC – mononuclear cells including lymphocytes and macrophages; PVC – perivascular cuff.

## References

1. Yao, H., et al. Molecular Architecture of the SARS-CoV-2 Virus. Cell 183, 730–738 e713 (2020).

2. Hu, B., Guo, H., Zhou, P. & Shi, Z.L. Characteristics of SARS-CoV-2 and COVID-19. Nat Rev Microbiol (2020).

3. Comar, C.E., et al. Antagonism of dsRNA-Induced Innate Immune Pathways by NS4a and NS4b Accessory Proteins during MERS Coronavirus Infection. mBio 10(2019).

4. Thornbrough, J.M., et al. Middle East Respiratory Syndrome Coronavirus NS4b Protein Inhibits Host RNase L Activation. mBio 7, e00258 (2016).

5. Nakagawa, K., Narayanan, K., Wada, M. & Makino, S. Inhibition of Stress Granule Formation by Middle East Respiratory Syndrome Coronavirus 4a Accessory Protein Facilitates Viral Translation, Leading to Efficient Virus Replication. J Virol 92(2018).

6. Niemeyer, D., et al. Middle East respiratory syndrome coronavirus accessory protein 4a is a type I interferon antagonist. J Virol 87, 12489–12495 (2013).

7. Rabouw, H.H., et al. Middle East Respiratory Coronavirus Accessory Protein 4a Inhibits PKR-Mediated Antiviral Stress Responses. PLoS Pathog 12, e1005982 (2016).

8. Xie, X., et al. An Infectious cDNA Clone of SARS-CoV-2. Cell Host Microbe 27, 841–848 e843 (2020).

9. Xie, X., et al. A nanoluciferase SARS-CoV-2 for rapid neutralization testing and screening of anti-infective drugs for COVID-19. Nat Commun 11, 5214 (2020).

10. Mulligan, M.J., et al. Phase I/II study of COVID-19 RNA vaccine BNT162b1 in adults. Nature 586, 589–593 (2020).

11. Hou, Y.J., et al. SARS-CoV-2 Reverse Genetics Reveals a Variable Infection Gradient in the Respiratory Tract. Cell 182, 1–18 (2020).

12. Thi Nhu Thao, T., et al. Rapid reconstruction of SARS-CoV-2 using a synthetic genomics platform. Nature 582, 561–565 (2020).

13. Xia, H., et al. Evasion of Type I Interferon by SARS-CoV-2. Cell Rep 33, 108234 (2020).

14. Kotaki, T., Xie, X., Shi, P.Y. & Kameoka, M. A PCR amplicon-based SARS-CoV-2 replicon for antiviral evaluation. Sci Rep 11, 2229 (2021).

15. Zhang, X., et al. A trans-complementation system for SARS-CoV-2 recapitulates authentic viral replication without virulence. Cell 184, 2229–2238 e2213 (2021).

16. Ricardo-Lax, I., et al. Replication and single-cycle delivery of SARS-CoV-2 replicons. Science 374, 1099–1106 (2021).

17. Ju, X., et al. A novel cell culture system modeling the SARS-CoV-2 life cycle. PLoS Pathog 17, e1009439 (2021).

18. Lohmann, V., et al. Replication of subgenomic hepatitis C virus RNAs in a hepatoma cell line. Science 285, 110–113 (1999).

19. Khromykh, A.A. & Westaway, E.G. Subgenomic replicons of the flavivirus Kunjin: construction and applications. J. Virol. 71, 1497–1505 (1997).

20. Shi, P.Y., Tilgner, M. & Lo, M.K. Construction and characterization of subgenomic replicons of New York strain of West Nile virus. Virology 296, 219–233 (2002).

21. Graham, R.L., Deming, D.J., Deming, M.E., Yount, B.L. & Baric, R.S. Evaluation of a recombination-resistant coronavirus as a broadly applicable, rapidly implementable vaccine platform. Commun Biol 1, 179 (2018).

22. Yount, B., Roberts, R.S., Lindesmith, L. & Baric, R.S. Rewiring the severe acute respiratory syndrome coronavirus (SARS-CoV) transcription circuit: engineering a recombination-resistant genome. Proc Natl Acad Sci U S A 103, 12546–12551 (2006).

23. Muruato, A.E., et al. A high-throughput neutralizing antibody assay for COVID-19 diagnosis and vaccine evaluation. Nat Commun 11, 4059 (2020).

24. Liu, Y., et al. The N501Y spike substitution enhances SARS-CoV-2 transmission. Nature, In press (2021).

25. Plante, J.A., et al. Spike mutation D614G alters SARS-CoV-2 fitness. Nature 592, 116–121 (2021).

26. Johnson, B.A., et al. Loss of furin cleavage site attenuates SARS-CoV-2 pathogenesis. Nature 591, 293–299 (2021).

27. Lau, S.Y., et al. Attenuated SARS-CoV-2 variants with deletions at the S1/S2 junction. Emerg Microbes Infect 9, 837–842 (2020).

28. Chen, R.E., et al. Resistance of SARS-CoV-2 variants to neutralization by monoclonal and serum-derived polyclonal antibodies. Nat Med 27, 717–726 (2021).

29. Muruato, A., et al. Mouse-adapted SARS-CoV-2 protects animals from lethal SARS-CoV challenge. PLoS Biol 19, e3001284 (2021).

30. Silvas, J.A., et al. Contribution of SARS-CoV-2 Accessory Proteins to Viral Pathogenicity in K18 Human ACE2 Transgenic Mice. J Virol 95, e0040221 (2021).

31. Kern, D.M., et al. Cryo-EM structure of SARS-CoV-2 ORF3a in lipid nanodiscs. Nat Struct Mol Biol 28, 573–582 (2021).

32. Freundt, E.C., et al. The open reading frame 3a protein of severe acute respiratory syndrome-associated coronavirus promotes membrane rearrangement and cell death. J Virol 84, 1097–1109 (2010).

33. Issa, E., Merhi, G., Panossian, B., Salloum, T. & Tokajian, S. SARS-CoV-2 and ORF3a: Nonsynonymous Mutations, Functional Domains, and Viral Pathogenesis. mSystems 5(2020).

34. Ren, Y., et al. The ORF3a protein of SARS-CoV-2 induces apoptosis in cells. Cell Mol Immunol 17, 881–883 (2020).

35. McAloose, D., et al. From People to Panthera: Natural SARS-CoV-2 Infection in Tigers and Lions at the Bronx Zoo. mBio 11(2020).

36. Kuchipudi, S.V., et al. Multiple spillovers from humans and onward transmission of SARS-CoV-2 in white-tailed deer. Proc Natl Acad Sci U S A 119(6):e2121644119., doi: 10.1073/pnas.2121644119 (2022).

37. Shi, J., et al. Susceptibility of ferrets, cats, dogs, and other domesticated animals to SARS-coronavirus 2. Science 368, 1016–1020 (2020).

38. Bonilla-Aldana, D.K., et al. SARS-CoV-2 natural infection in animals: a systematic review of studies and case reports and series. Vet Q 41, 250–267 (2021).

39. Valencak, T.G., et al. Animal reservoirs of SARS-CoV-2: calculable COVID-19 risk for older adults from animal to human transmission. Geroscience 43, 2305–2320 (2021).

40. Pomorska-Mol, M., Wlodarek, J., Gogulski, M. & Rybska, M. Review: SARS-CoV-2 infection in farmed minks - an overview of current knowledge on occurrence, disease and epidemiology. Animal 15, 100272 (2021).

41. Liu, Y., et al. Neutralizing Activity of BNT162b2-Elicited Serum. N Engl J Med 384, 1466–1468 (2021).

42. Liu, Y., et al. BNT162b2-Elicited Neutralization against New SARS-CoV-2 Spike Variants. N Engl J Med 385, 472–474 (2021).

43. Liu, J., et al. BNT162b2-elicited neutralization of B.1.617 and other SARS-CoV-2 variants. Nature (2021).

44. Menachery, V.D., et al. Attenuation and restoration of severe acute respiratory syndrome coronavirus mutant lacking 2’-o-methyltransferase activity. J Virol 88, 4251–4264 (2014).

45. Kee, J., et al. SARS-CoV-2 ORF8 encoded protein contains a histone mimic, disrupts chromatin regulation, and enhances replication. bioRxiv, doi: https://doi.org/10.1101/2021.1111.1110.468057 (2022).

46. Su, Y.C.F., et al. Discovery and Genomic Characterization of a 382-Nucleotide Deletion in ORF7b and ORF8 during the Early Evolution of SARS-CoV-2. mBio 11(2020).

47. Young, B.E., et al. Effects of a major deletion in the SARS-CoV-2 genome on the severity of infection and the inflammatory response: an observational cohort study. Lancet 396, 603–611 (2020).

48. Xie, X., et al. Engineering SARS-CoV-2 using a reverse genetic system. Nature Protocols 16, 1761–1784 (2021).

49. Ku, Z., et al. Molecular determinants and mechanism for antibody cocktail preventing SARS-CoV-2 escape. Nature Communications, https://doi.org/10.1038/s41467-41020-20789-41467 (2021).

